# Functional redundancy and anaerobic metabolism characterize the skin microbiome of a critically endangered sawfish

**DOI:** 10.64898/2026.03.05.709720

**Authors:** Michael P. Doane, Belinda Martin, Emma Kerr, Elizabeth A. Dinsdale, Malak Malak Rangers, Leonardo Guida, Pete M. Kyne

## Abstract

Using shotgun metagenomics of largetooth sawfish (Pristis pristis) skin and paired pool water samples, we provide the first taxonomic and functional characterization of a sawfish skin microbiome and assess its ecological distinctiveness, assembly, and resilience relative to the surrounding water column. The skin microbiome was strongly host-associated and compositionally distinct, with lower taxonomic richness than the water column but dominated by Bacillota, particularly spore-forming Bacilli. Functional profiling revealed enrichment of genes associated with sporulation, dormancy, anaerobic metabolism, and peptide transport, consistent with adaptation to low-oxygen, low-flow pool conditions. In contrast, water-column microbiomes were more taxonomically diverse and enriched in phototrophic and polysaccharide-utilization pathways. Despite reduced taxonomic diversity, sawfish skin communities exhibited higher functional redundancy, with gene functions accumulating more rapidly per taxon. This pattern supports a host-filtered, lottery-like assembly process that produces taxonomically variable yet functionally conserved communities. The enrichment of dormancy and anaerobic pathways suggests the skin microbiome persists through periods of host quiescence and environmental stress while maintaining metabolic potential. Together, these results demonstrate that the sawfish skin supports a resilient, functionally robust microbiome distinct from ambient aquatic communities, highlighting the potential for integrating microbial functional data into conservation strategies for Critically Endangered vertebrates.

## Introduction

Globally, animal and plant species face unprecedented conservation challenges as human activities instigate widespread alterations to natural environments, leading to biodiversity decline (Keck *et al*., 2025). Despite our best efforts, many protected species, including large vertebrates, continue to decline (McCauley *et al*., 2015; Darwall *et al*., 2018), suggesting unseen biological processes are failing well before impacts are observed. Increasingly, the microbiome has emerged as a critical determinant of host resilience that influences health, adaptability, and survival amid habitat degradation, climate shifts, and biodiversity loss (Cavicchioli *et al*., 2019). The microbiome is now recognized as a fundamental feature of all animals, with microbial community composition structured non-randomly and performing essential functions that maintain host health and fitness. Disruption in the relationship between the host and its microbiomes corresponds to cascading species impacts (Bordenstein *et al*., 2024). For species of conservation concern, understanding how a normal microbiome assembles with the host is critically important towards advancing conservation measures (Wu *et al*., 2024).

Intervention in the form of microbiome modulation has been proposed in some instances to counter the effects of these impacts, such as climate change, to make corals more resilient to increased temperatures (Peixoto *et al*., 2022). While such reactive efforts to ‘force’ adaptive responses are underway, there are, however, critical knowledge gaps around basic underlying ecological processing shaping the host microbiome (Walling *et al*., 2025). This gap is particularly pronounced for non-model, threatened species such as elasmobranchs (sharks and rays), whose skin microbiomes remain poorly characterized despite their potential importance for immune defence and adaptation to fluctuating aquatic environments (Doane *et al*., 2017; Apprill *et al*., 2020). Framing these microbiomes as dynamic ecosystems in their own right, governed by ecological processes such as selection, drift, dispersal, and diversification, offers a powerful means to uncover how host-associated communities contribute to resilience and health. Central to the conservation efforts is the preservation of function, with ecological processes of community assembly forming the foundational scaffold. Because large-bodied elasmobranchs in estuarine systems are disproportionately affected by habitat degradation, resolving the ecological processes and functional potential of their skin microbiomes represents a necessary step toward conservation strategies that prioritize the preservation of host-associated function.

Sawfish (family Pristidae) are among the most severely impacted elasmobranchs, with all five species now assessed as Critically Endangered on the IUCN Red List of Threatened Species (IUCN, 2025). Sawfish inhabit freshwater, estuarine, and coastal habitats. The central threats to these species include entanglement in fishing gear, overexploitation, and habitat degradation (Espinoza *et al*., 2022). Habitat alteration includes development and water diversions for agriculture. Degradation of freshwater environments, in particular, has driven steep declines in water quality, with less than one-fifth of preindustrial freshwater wetlands remaining (Albert *et al*., 2021; Ross and Randhir, 2022), resulting in significant biodiversity loss, specifically for vertebrate species (Reid *et al*., 2019). For species such as the largetooth sawfish (*Pristis pristis*), which rely on freshwater habitats as nursery areas, water quality is critical to their survival. Within northern Australia, largetooth sawfish occur in river systems that experience dynamic seasonal wet-dry cycles, often sequestering them to isolated pools until monsoonal rains occur. During this time, the body condition of sawfish can diminish rapidly (Lear *et al*., 2021), signalling stress to the animal. These periods of stress are expected to slow metabolic function, leaving these animals susceptible to their environment.

The microbiome is a critical modulator of host health, playing an instrumental role in shaping the physiological, immune, and ecological functions essential to the host’s survival and well-being (Opstal and Bordenstein, 2015). Among host-associated microbiomes, the skin represents a critical ecological interface between the organism and its environment. The microbiome mediates pathogen exclusion, chemical communication (Turnbaugh *et al*., 2007; Sylvain *et al*., 2020), and in aquatic species, host-microbial biofilms reflect both host physiology and water quality (Doane *et al*., 2022; Wilde *et al*., 2024). The microbiomes’ integrative role in health, especially for non-human model species, is still emerging. Marine fish species microbiomes are diverse, displaying host specificity (Doane *et al*., 2017); however, they are also adaptable, adjusting to environmental conditions (Chiarello *et al*., 2018; Sylvain *et al*., 2020). For instance, the microbiome of 73 whale sharks (*Rhincodon typus*) collected across 5 locations around the world showed similar microbial network properties and several metagenome-assembled genomes were present on sharks from the same and across locations (Doane *et al*., 2023b). In contrast, 13 sympatric fish species from the Mediterranean Sea harbor distinct microbiome communities (Ruiz-Rodríguez *et al*., 2020). A review of 36 studies characterizing the microbiome of 98 fish species shows that physicochemical properties drive microbiome composition (Bell *et al*., 2024). The collective interplay of the host and the environment in shaping the microbiome signifies its adaptive capacity, potentially acting analogously to epigenetic phenotypical adjustments (Anka *et al*., 2024). In static, low-oxygen pools characteristic of the northern Australian dry season, microbial communities on the sawfish skin likely experience the same environmental constraints as their host. As such, persistence may depend on physiological strategies like dormancy, sporulation, and anaerobic energy metabolism—traits that maintain microbial stability during periods of host quiescence.

When a host enters dormancy or a slowed metabolic state, the microbiome can restructure to meet physiological demands (Brown *et al*., 2022). For instance, the gut microbiome of the thirteen-lined ground squirrel (*Ictidomys tridecemlineatus*) shifts to support nitrogen recycling during periods of fasting (Regan *et al*., 2022). The microbiome also serves as a critical guardian for its host during periods of slowed metabolism. For example, while the skin microbiome is predicted to influence susceptibility to *Pseudogymnoascus destructans*, the causative agent of white-nose syndrome in Nearctic bats, Palearctic bats tolerate high burdens of *P. destructans* without pathology, a resilience likely attributed to antifungal activity within their skin microbiome (Troitsky *et al*., 2023). For animals such as the sawfish, isolation in stagnant pools coupled with slowed metabolic states may increase susceptibility to environmental exposure. Therefore, as an extension of the sawfish’s immune response, we predict the microbiome will be highly structured and distinct from the surrounding water column, reflecting selective filtering by the host. For example, the healthy water column is typically dominated by free-living, phototrophic carbon fixers adapted to planktonic life (Doane *et al*., 2023a), whereas host-associated microbiomes are enriched for taxa with biofilm-forming and surface-colonization capacities, as observed in leopard sharks (Goodman *et al*., 2022). Such functional partitioning highlights the transition from a free-living to a host-selected microbial assemblage.

While microbiome taxonomic composition reflects “who is there,” functional attributes reveal “what they are doing” (Dinsdale *et al*., 2008), and therefore how microbial communities can persist and adapt under environmental stress. Functional gene composition provides critical insight into the ecological roles and resilience mechanisms of host-associated microbiomes. Many microbiome studies rely on 16S rRNA amplicon sequencing to infer community structure and often use predictive tools such as PICRUSt or Piphillin to estimate functional potential. However, these predictive approaches frequently fail to capture the true functional repertoire of microbial communities, particularly in novel or under-characterized environments, where taxa-function relationships are less constrained (Louca *et al*., 2016). In such contexts, relying solely on taxonomy misrepresents the ecological capabilities of the microbiome, leading to incomplete or inaccurate interpretations. Directly measuring gene functions through metagenomic sequencing is therefore essential for identifying the metabolic and ecological traits that sustain microbial communities, particularly when assessing how microbiomes contribute to host health and resilience. Understanding the functional capacity of microbiomes can inform management decisions, such as predicting host responses to environmental stressors or designing microbiome-based interventions to enhance species persistence and is critical for successful conservation.

Together, these observations suggest that the sawfish skin supports a structured microbial community with unique metabolic capacities relative to the ambient environment. We hypothesize that host-driven filtering selects for specific gene functions, resulting in a microbiome that is functionally adapted to its host. To test this hypothesis, we aimed to: (1) describe the taxonomic composition and functional potential of the sawfish skin microbiome, (2) determine its connectivity and distinctiveness relative to the surrounding water column microbiome, and (3) identify ecological assembly patterns and key microbial functions that underpin microbiome resilience under low-oxygen, low-flow conditions. By integrating taxonomic and functional profiles, we reveal a highly selective microbiome structured for persistence under oxygen limitation, suggesting that host-driven filtering promotes community stability under environmental stress.

## 1. Experimental Procedures

### 2.1 Sampling location and environmental data collection

Largetooth sawfish (*Pristis pristis;* hereafter ‘sawfish’) were caught and sampled on the Daly River floodplain in the Northern Territory of Australia, ∼130 km southwest of Darwin (Figure 1). Nine sawfish were sampled from an isolated floodplain pool during the late dry season when these waterholes are disconnected from the perennially flowing main river channel. Sample collection occurred on three separate days in August-September 2021. Sawfish ranged in size 99.6-120.0 cm total length and were all aged 0+ (i.e., young-of-the-year) (Supplemental Table S1). Sawfish were sampled as part of a long-term ‘rescue’ program whereby sawfish stranded in rapidly drying waterholes are relocated to the permanent water of the main river channel in order to enhance survivorship of this Critically Endangered species. Water was also collected on each sampling day in triplicate (1.5 L). Environmental data collected included date and time along with water temperature (°C), specific conductivity (mS/cm), dissolved oxygen (mg/L and %), salinity (PSU), and turbidity (NTU). For sawfish, the sex, total length (mm), and capture depth (m) were recorded. Sawfish were sampled from a freshwater (salinity: 0.13-0.15 PSU) pool over several days that varied from mildly hypoxic to hypoxic (63.5-36.6 % saturation).

**Figure 1:**
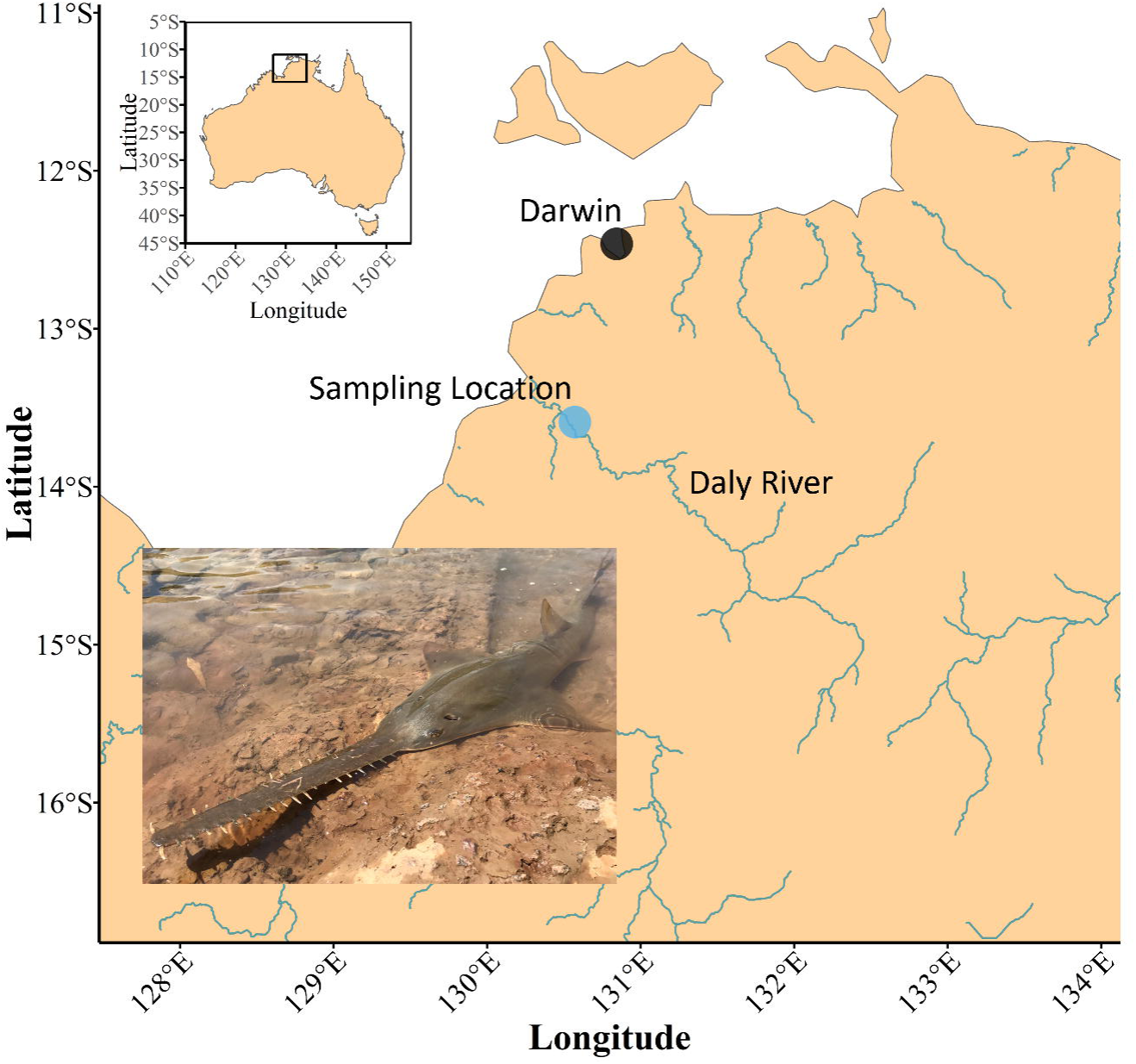
Map of the sampling location for largetooth sawfish (*Pristis pristis*) in the Northern Territory, Australia. The photo inset shows a largetooth sawfish. The black dot indicates the nearest city, Darwin, Northern Territory, and the blue dot marks the pool where individual sawfish were sampled. The Daly River and its tributaries are labeled. Photo credit: Pete Kyne.

### 2.2 Sampling collection and DNA extraction

Microbial samples were collected using nylon flocked swabs (Regular FLOQSwab®, Copan, CA, USA) applied along the dorsal surface beneath the first dorsal fin. Swab tips were aseptically broken into 1.5 mL tubes and sealed. Water column samples were collected at the three timepoints by filtering 1.5 L of seawater onto 0.45 µm Sterivex™ filters. All samples were stored on ice before transfer to −20°C storage. Samples were shipped to Flinders University (Adelaide, Australia), where DNA was extracted and prepared for metagenomic sequencing.

For swabs, the NucleoSpin Tissue kit (Machery-Nagel, Germany) was used for extraction and DNA purification, following manufacturer recommendations with slight modifications in reagent volumes, as outlined in Martin *et al*., (2024). First, samples were thawed at room temperature. Swabs were then placed in 5 mL tubes for pre-lysis in 270 µL T1 buffer with 37.5 µL of Proteinase K (final concentration of 2.5 mg/mL) and incubated at 55 °C overnight with rotation. Lysis was then performed in 300 µL of B3 buffer with incubation for 10 minutes at 70 °C with rotation. The swab was removed, and a DNA IQ™ spin basket (Promega Corporation) was placed into the tube containing the lysed material. The swab was then placed into the spin basket and spun at 14,000 rpm for 60 seconds to remove all material from the swab. The swab and basket were then discarded. DNA binding conditions were then adjusted to capture DNA onto silica filter members with 300 µL of 100 % ethanol. Remaining steps followed the manufacturer’s standard protocol with DNA eluted off the filter with 50 µL of BE heated to 70 °C, with incubation of 10 minutes before spinning at 14,000 rpm for 60 seconds. This was repeated to bring the total eluted volume to 100 µL. DNA concentrations were quantified with the Qubit 4-Fluorometer using the HS dsDNA kit (Invitrogen, USA) and then stored at −20 °C until further processing.

Water column DNA was extracted and purified with the NucleoSpin Tissue kit following the methods outlined in Doane et al., (2023). Briefly, 720 µL of T1 with 90 µL of Proteinase K (final concentration 2.25 mg/mL) was placed into the Sterivex™ and incubated overnight. The solution was suctioned from the Sterivex™ using a 5 ml luer lock syringe and placed into a 5 mL Eppendorf tube, with 800 µL B3 for subsequent lysis. Binding conditions were adjusted using 800 µL 100% ethanol, and successive steps followed the standard manufacturer’s protocol, as outlined above. DNA concentrations were quantified with the Qubit 4-Fluorometer using the HS dsDNA kit and then stored at −20 °C until further processing. DNA-negative controls were extracted in parallel for the swabs and Sterivex™ to ensure no reagent contamination.

### 2.3 DNA libraries, MGI sequencing, and data quality control

The samples were then prepared into shotgun metagenomic libraries using the xGen 2S library prep kit (Integrated DNA Technologies, USA). The DNA was first mechanically sheared into fragments of approximately 450 bp (2 x 150) or 650 bp (2 x 300) using the Q800R sonicator (QSONICA, USA). For 450 bp fragments, 50 µL of purified DNA was placed into 0.3 mL thin-walled PCR tubes and sonicated for 9 minutes at 15 seconds on and 15 seconds off with 20 % amplitude. For 650 bp fragments, similar volumes were sonicated for 6 minutes with 15 seconds on and 15 seconds off at 20 % amplitude. The sheared DNA was then taken forward for library prep per manufacturer recommendations, following the ‘less than 10ng gDNA’ methods. The DNA first underwent end repair, and adaptors were ligated to each end. An indexing PCR was then performed using 12 cycles with manufacturer cycle temperatures, followed by a PCR cleanup. Libraries were sent to the South Australian Genomics Center for sequencing with the MGI DNBSEQ G400 (MGI, China). Two sequencing runs were conducted at 2 x 150 bp and 2 x 300 bp lengths. The libraries were prepared using the MGIEasy kit (MGI, China) to convert linear DNA libraries into the MGI DNA nanoball structures. Rolling circle amplification was then performed, yielding a nanoball that is composed of repeating regions of the original sequence. These are then loaded onto the flow-cell for sequencing. Two sequencing runs were performed. The initial 2 × 150 bp run yielded incomplete library recovery, with several samples failing to generate libraries. Consequently, a second sequencing run was conducted using 2 × 300 bp chemistry following the phase-out of the 2 × 150 bp configuration on the MGI DNBSEQ-G400 platform.

### 2.4 Sequencing data analysis

The resulting fastq files were processed using the ATAVIDE_LITE pipeline for downstream analysis (Edwards, 2025)(https://github.com/linsalrob/atavide_lite). The pipeline first implements Fastp (Chen, 2023) to remove sequencing adapters and discard reads with more than one ambiguous base or shorter than 100 bp. The host DNA was then estimated and removed by mapping reads with minimap2 (Li, 2018) to the smalltooth sawfish (*Pristis pectinata*) genome (Genbank: GCF_009764475.1). The taxonomic and gene function read assignments were then made using MMseqs2 (Hauser et al., 2016). Reads were compared against the UniRef50 database (March 2023) and assigned UniRef IDs. Taxonomic assignments were performed using a lowest common ancestor (LCA) approach based on UniRef50 clustering (>50% identity). Each sequence was assigned to a UniRef cluster and annotated to the most specific shared taxonomic rank among matching proteins. When matches spanned multiple taxa, classification was conservatively resolved to their lowest shared rank. For example, if a read matched genomes within the phylum Bacilliota belonging to different families, it was classified as Family_unknown. Taxonomic hierarchical organization was then reformatted with taxonkit (v0.20.0)(Shen and Ren, 2021). Gene function UniProtIDs were mapped to proteins associated with BV-BRC subsystems (Olson *et al*., 2023) and formatted in the SEED Subsystem hierarchical organization of nested functional categories, progressing from broad groupings (Level 1) to increasingly specific functional assignments (Levels 2, 3, and Subsystem). This generated two data frames summarizing taxonomic and functional annotations, with each entry representing the number of reads per sample assigned to a given taxon or gene function. Data processing was done on the Flinders University HPC DeepThought (Flinders University High-Performance Computing cluster, 2021).

Microbial species representing less than 0.0001% of total reads were considered spurious assignments and removed. Species-level counts were then collapsed to the Family level by summing abundances, and all subsequent analyses were performed on either count or proportional data (as noted below). No reads were removed from the potential gene function data set. Gene functions were standardized to reads per million (RPM) rather than proportional abundance, as is typical for taxonomic datasets, because functional read counts can include fractional values. This occurs during the annotation process when a sequence matches multiple protein-coding genes equally, with the read count divided among those matches. Standardizing to RPM preserves these fractional contributions and allows for meaningful comparisons of gene function abundances across samples, ensuring that multi-matching reads are appropriately accounted for.

### 2.5 Community data analysis

Microbial communities were compared across the sawfish skin (environment: *skin*) and the surrounding water (environment: *water*). All analyses of the taxonomic dataset was performed at the Family classification. Alpha and gamma diversity were calculated as richness for the two environments using *specnumber* and *diversity* functions in vegan (v2.6.4)(Oksanen *et al*., 2020). Statistical differences in alpha diversity patterns between environments was calculated using the Dunn test in *Dunn.test* function in dunn.test package (version 1.3.6) (Dinno, 2014). For community analysis, both datasets (taxonomy: proportion; gene functions: RPM) were first fourth-root transformed, and beta-diversity was calculated as a Bray-Curtis dissimilarity matrix and ordinated using the *metaMDS* function in vegan (v2.6.4) with autotransform = FALSE. Taxa and gene function composition was compared between environments using the *adonis2* function in vegan (9999 permutations)(Anderson, 2001). Enriched taxonomic and gene function groups between environments were identified using the Analysis of Compositions of Microbiomes with Bias Correction (ANCOM-BC) method implemented in the ANCOM-BC R package (v1.6.4)(Lin and Peddada, 2020). The analysis was performed using count data (gene functions = RPM). Datasets were converted to phyloseq objects using the phyloseq package in R (v1.46.0) (McMurdie and Holmes, 2013), and subsequently to TreeSummarizedExperiment (*tse)* objects using the *makeTreeSummarizedExperimentFromPhyloseq* function from the mia package (v1.10.0)(Ernst *et al*., 2023). The *tse* object is a hierarchical data structure that stores abundance matrices alongside sample metadata, taxonomic annotations, and phylogenetic relationships, providing the format required for ANCOM-BC analyses. ANCOM-BC was run with the environment factor as the main explanatory variable. Parameters included a prevalence cutoff of 0.10 (prv_cut = 0.10), library size threshold of 1,000 reads (lib_cut = 1000), Holm correction for multiple testing, and structural zero detection enabled. Enriched (features identified as differentially abundant) taxa and gene functions were correlated together using Spearman’s rank correlation, and only taxa was clustered using *hclust* complete linkage. Heatmap visualization and clusters was done using the pheatmap (v1.0.13)(Kolde, 2025). The relationship between taxa richness and predicted gene functions was modeled using the Michaelis–Menten equation implemented via nonlinear least squares fitting (base R::*nls*). The model estimated parameters describing the saturation curve and the half-saturation constant (K). To quantify the overall accumulation of gene functions, the area under the curve (AUC) was calculated using the trapezoidal integration method using *trapz*() function in pracma package (v2.4.6)(Borchers, 2025) applied to the fitted curves. These analyses provided complementary measures of functional diversity efficiency (K) and total capacity (AUC) for each microbiome type. All visualizations, unless noted, were generated in ggplot2 (v4.0.0) (Wickham, 2016). All analyses were conducted in R (v4.5.1). All scripts can be found at https://github.com/mpdoane2/Sawfish-microbiome-analysis.

## 2. Results

### 3.1 Shotgun metagenomic libraries

We analyzed 14 shotgun metagenomic libraries from sawfish skin (n = 9) and the bacterioplankton in the pool water (n = 5). The mean raw number of sequences for sawfish was 7,837,125 (standard error ± 2,086,015) and water was 9,992,099 (± 1,990,651). After host filtering, the mean number of reads was 3,580,491 (± 1,279,530) and 9,914,000 (± 2,222,261), which accounted for approximately 45.7 % and 98.0 % of quality-controlled reads from sawfish and water, respectively. From the host-filtered reads, 22.2 % (± 1.1 %) and 30.0 % (± 1.4 %) of sawfish reads were assigned a taxonomy and gene function, respectively, while 16.2 % (± 0.3 %) and 23.2 % (± 0.8 %) of water reads were assigned a taxonomy and gene function, respectively (Supplemental Fig S1; Supplemental Table S1). Therefore, many microbes in the water and on the sawfish remain uncharacterised.

### 3.2 Distinct taxonomic microbiomes from sawfish and water column communities

The sawfish and water column microbiomes were highly distinctive. The most abundant taxa classified to the phylum level (excluding ‘Bacteria unknown’) were Bacillota (sawfish: 56.6 %; water: 30.1 %), Pseudomonadota (4.48 %; 14.91 %), and Actinomycetoa (3.17 %; 9.42 %). At the family level, the top five most abundant groups for sawfish included the family Bacillaceae (25.6 ± SE 1.25 %), order Bacillales (19.4 % ± 1.37 %), phylum Bacillota (3.77 ± 0.21 %), class Bacilli (2.70 ± 0.15 %), and family Paenibacillacea (1.67 ± 0.25 %). The most abundant microbial family groups for the water bacteriaplankton included the order Bacillales (13.5 ± 1.22 %), family Bacillaceae (11.1 ± 1.24 %), class Actinomycetes (4.14 ± 0.19 %), phylum Pseudomonadota (3.30 ± 0.42 %), and phylum Planctomycetota (2.21 ± 0.33 %) (Fig 2A; Supplemental Table S2).

**Figure 2:**
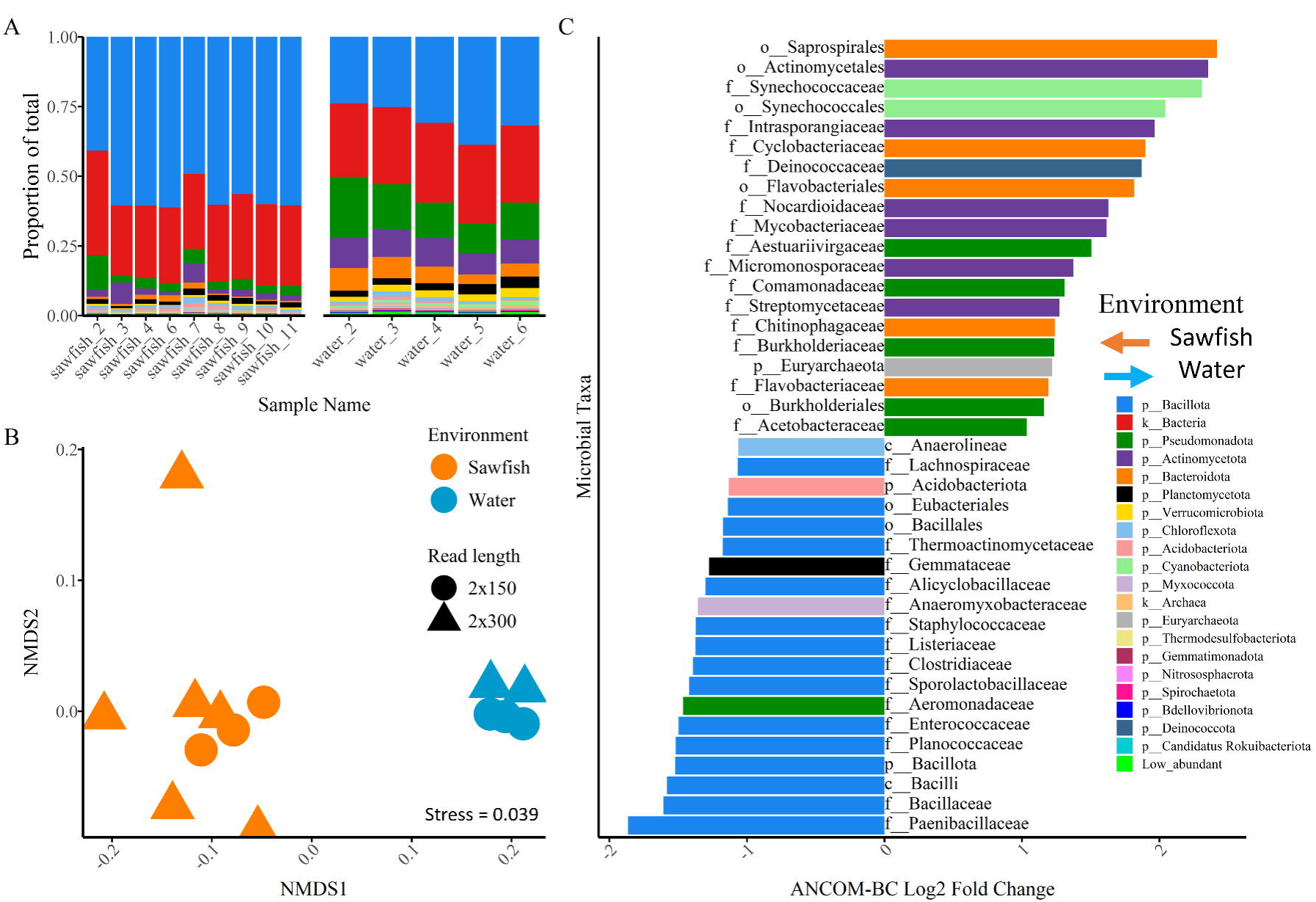
Microbial composition and richness of largetooth sawfish (*Pristis pristis*) skin and surrounding water from the Daly River floodplain, Northern Territory, Australia. **(A)** Relative abundance of microbial taxa assigned at the phylum level. The 20 most abundant phyla are shown; remaining taxa are grouped as *Low_abundant*. Taxonomic assignments were made using a least common ancestor approach. Therefore, reads classified only to Kingdom Bacteria or Archaea represent sequences matched to multiple phyla within those respective kingdoms (see *2.5 Community data analysis* section. The phylum legend is shared with panel C. **(B)** Ordination of microbial families based on Bray Curtis similarity. Point color corresponds to sample environment (sawfish and water) and point shape indicates sequencing read length. **(C)** Log fold change (LFC) abundance distribution for the top 20 microbial families across the two microbiome environments (sawfish and water). Right extending bars represent families enriched in the water column, whereas left extending bars represent families enriched in the sawfish microbiome. Log fold change is shown along the x axis, where 1 (−1) corresponds to a two-fold difference in abundance in the direction of the bar. Microbial family names are displayed along the y axis emerging from zero in the opposite direction of each bar. Bar colors correspond to the phylum to which each family belongs and are consistent with panel A.

A total of 1143 families were identified in the combined sawfish and water column microbial dataset. Of those, 15 families were unique to the sawfish microbiome (1.3 % of total family richness), and 148 families were unique to the water column (12.9 %). The remaining 980 families were shared among the two environments, accounting for 85.7 % of all families found (Supplemental Fig S2A & Supplemental Fig S3). A similar trend is observed at the genus level, with 3.2% of genera unique to sawfish, 26.4 % to the water, and 70.3 % shared among the two environments (Supplemental Fig S2B). The total number of families identified for sawfish was 995, and 1128 for the water microbiome (Supplemental Fig S2A). The mean number of families identified in the sawfish microbiome was significantly less (895.0 ± 36.3; Kruskal-Wallis X^2^ = 9.04, p < 0.001) than the water microbiome (1122.4 ± 3.67; Supplemental Fig S3). The Shannon diversity at the family level of the sawfish microbiome was also significantly less than the water column (Kruskal-Wallis X^2^ = 9.00, p < 0.001; Supplemental Fig S4).

The composition of the microbiome was compared across the two environments. The microbiomes were significantly different in their family level composition, forming distinct patterns in ordination (PERMANOVA: pseudo-F (1,12) = 19.85, R^2^ = 0.623, p < 0.001; Fig 2B; Supplemental Fig S5). Day was included as a factor in the ordination to assess temporal variation. Variation among samples from both environments across days was minimal relative to the effect of environment (sawfish vs. water). Day was not quantitatively tested due to limited replication within sampling day (Supplemental Fig S6). Retaining all microbial families with a two-fold or greater abundance difference (LFC ≥ 1/−1) resulted in 284 significantly enriched families from 59 phyla across two kingdoms (Fig 2C; Supplemental Table S2).From the sawfish, 123 enriched microbial families were from Bacteria and four from Archaea. The top five enriched Bacterial groups included Bacillaceae family (Bacillota; q < 0.001), Bacillales order (Bacillota; q < 0.001), Bacillota phylum (Bacillota; q < 0.001), Bacilli class (Bacillota; q < 0.001), and Paenibacillaceae family (Bacillota; q < 0.001). The only Archaea from the skin microbiome of sawfish near 0.05% abundance was Methanocellacea family (Euryarchaeota; q < 0.001). From the water column, 149 out of 157 enriched microbial families were from Bacteria and eight from Archaea. The top five most abundant Bacterial families included those from Micromonosporaceae family (Actinomycetota; q < 0.001), Mycobacteriaceae family (Actinomycetota; q < 0.001), Burkholderiales order (Pseudomonadota; q < 0.001), Burkholderiaceae family (Pseudomonadota; q < 0.001), and Chitinophagaceae family (Bacteroidia; q < 0.001). From Archaea the two most abundant families greater than 0.05 % included Euryarchaeota phylum (Euryarchaeota; q < 0.001), and Candidatus Woesearchaeota phylum (Candidatus Woesearchaeota; q < 0.001).

### 3.3 Gene function variation among sawfish and water microbiomes

The microbiome community’s gene functions were investigated to determine if the host skin niche, like the taxonomic profiles, were distinct from the water column environment. A total of 774 SEED sub-systems were identified across the whole dataset, with 763 for sawfish and 766 for the water column. A similar number of Level 1 pathways and their corresponding abundances were identified across the two environments (Supplemental Fig S7 & S8). The most subsystem-rich Level 1 group for both environments were metabolism (sawfish: 236.3 ± 4.25 subsystems; water: 250.0 ± 3.04), followed by stress response, defence, and virulence subsystems (93.0 ± 1.52; 84.78 ± 3.27), and energy subsystems (87.8 ± 12; 79.56 ± 2.49). At the subsystem levels, the gene functions were distinct among the two environments (PERMANOVA: Pseudo-F (1,12) = 9.31, R^2^ = 0.44, p < 0.001; Fig 3A; SI Fig S9). Of the 774 subsystems, 256 differed significantly based on differential abundances (DA) analysis, with 101 subsystems having a 2-fold abundance difference (LFC ≥ 1/−1; SI Table S3). For sawfish, the top five DA subsystems (filtered LFC > 1 and abundance ranked; Fig 3B) were all involved in sporulation, including sporulation gene orphans (q < 0.01), spore germination (q < 0.01), sporulation proteins SpoVA cluster (q < 0.05), spore germinant receptors (q < 0.001), and sporulation proteins SigEG cluster (q < 0.05). Considering the top abundant and DA subsystems (including LFC < 1 and abundance ranked) Dpp dipeptide ABC transport system (Level 1: Membrane transport; q < 0.001), flagellum (Cellular processes; q < 0.001), type IV pilus (Membrane transport; q < 0.01), and lysine fermentation (Metabolism; q < 0.05) were enriched in the sawfish. In the water column, functions with the greatest DA (filtered LFC > 1 and abundance ranked) included utilization systems for glycans and polysaccharides (Membrane transport; q < 0.001), chlorophyll biosynthesis (Metabolism; q < 0.001), biosynthesis of arabinogalactan in mycobacteria (Cell envelope; q < 0.001), proteorhodopsin (Energy; q < 0.001), and photosystem II (Energy; q < 0.001). When considering the top five abundant DA subsystems (including LFC < 1 and abundance ranked) from water gram-negative cluster (Cell envelope; q < 0.01), and several functions from Protein processing Level 1 groups including ribosomal proteins, single copy (q < 0.01), chaperones GroEL/GroES and thermosome (q <0.0001), protein chaperones (q < 0.0001), and translation elongation factors (q < 0.0001).

**Figure 3:**
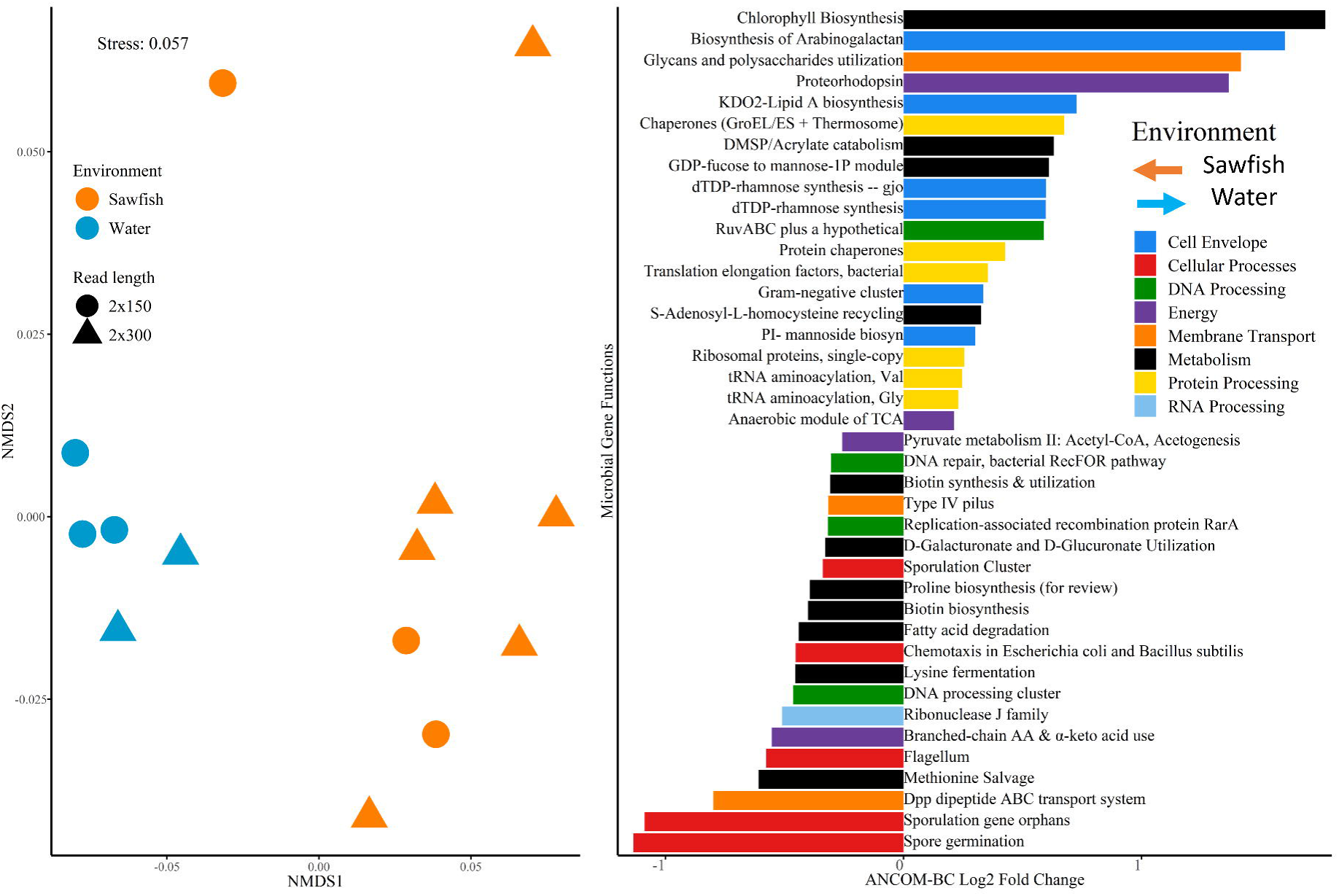
Gene function composition across the sawfish microbiome and the surrounding water column. **(A)** Nonmetric multidimensional scaling (nMDS) ordination of gene function profiles from sawfish and water microbiomes. Samples cluster distinctly by environment, with high similarity among replicate samples. Distances between points reflect relative similarity in gene function composition, calculated using Bray Curtis similarity on fourth root transformed reads per million normalized data. Blue denotes water column samples and orange denotes sawfish samples. Circles represent 2 × 150 bp libraries and triangles represent 2 × 300 bp libraries. **(B)** Log fold change (LFC) abundance distribution of differentially abundant microbial Subsystem functional groups between sawfish and water microbiomes. Right extending bars (positive LFC) represent Subsystems enriched in the water column, whereas left extending bars (negative LFC) represent Subsystems enriched in the sawfish microbiome. Log fold change is shown along the x axis, where 1 (−1) corresponds to a two-fold increase in abundance in the direction of the bar. Subsystem names are displayed along the y axis emerging from zero in the opposite direction of each bar. Bar colors correspond to the Level 1 functional category to which each Subsystem belongs.

The composition of subsystems within Energy was also significantly different among the two environments (PERMANOVA: pseudo-F (1,12) = 11.39, R2 = 0.49, p < 0.001; SI Fig S10). There were seven DA energy subsystems enriched in the sawfish including pyruvate metabolism II (q < 0.01; SI Table 3), branched-chain amino acids and alpha-keto acids utilization (q < 0.001), fermentations: mixed acids (q < 0.01), cytochrome d ubiquinol oxidase operon (q < 0.001), pyruvate formate-lyase cluster (q < 0.001) and electron-bifurcating caffeyl-CoA reductase-Etf complex (q < 0.001). The top five from the water included anaerobic module of TCA (q < 0.01), proteorhodopsin (q < 0.001), photosystem II (q < 0.001), acetoin, butanediol metabolism (q < 0.001), ethylmalonyl-CoA pathway of acetyl-CoA assimilation (q < 0.001).

### 3.4 Contrasting gene function redundancy from sawfish and the water column microbiome

The differentially abundant taxa and functions were compared to identify taxon-function correlation (Fig 4). Correlative trends reveal distinct relationships between taxa and gene functions, with two taxonomic groups (as evidenced by clustering) that align with the DA analysis. A similar trend is observed when considering all taxa and all gene functions, thereby demonstrating results are ecological rather than methodological (SI Fig S11). This indicates that taxa found in similar environments share similar gene functions. Cluster one was composed of 82 taxa with significant DA in water and six with DA significant to the sawfish. Cluster two contained 50 DA taxa, all from sawfish (from here on Cluster 1 referred to as water cluster and Cluster 2 is sawfish cluster). In the sawfish cluster, two groups emerged, one composed primarily of families from the phylum Bacillota, with the second group composed of ‘other’ microbial phyla. This second group formed a more uniform correlation pattern with relatively weak correlation across the gene functions.

**Figure 4:**
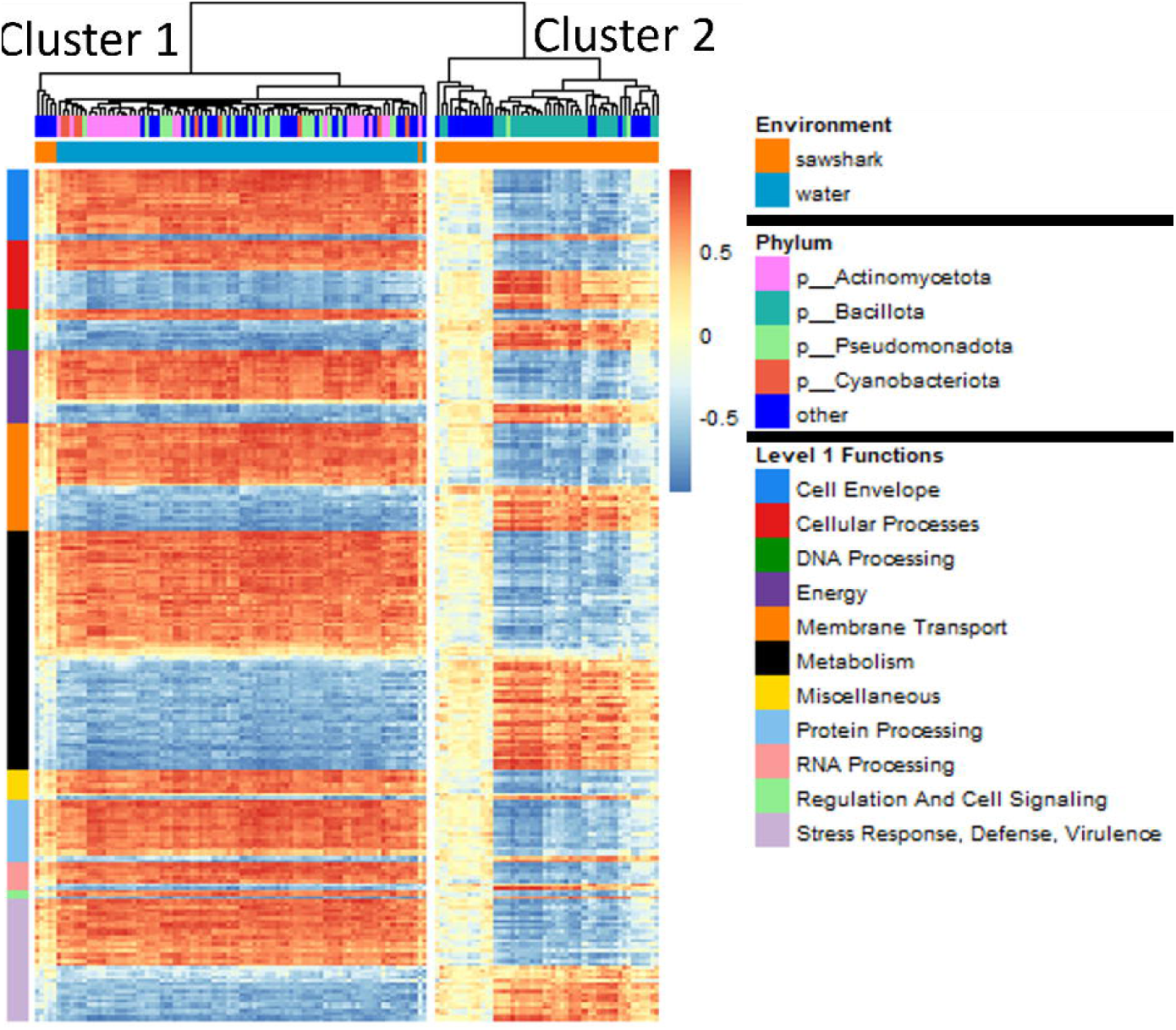
Heatmap illustrating relationships between significantly differentially abundant (DA) microbial taxa (columns) and gene functions (rows) in the sawfish (orange) and water column (blue) microbiomes. Taxa with a mean relative abundance < 0.001 were excluded prior to heatmap construction. Colors represent Spearman correlation coefficients, where red indicates positive correlation and blue indicates negative correlation; lighter shades denote weaker associations. Columns correspond to microbial families, labeled by their phylum classification and the environment in which they are significantly enriched. Rows correspond to gene functions, organized by mean abundance across the dataset within their Level 1 SEED category (broadest functional classification).

Two groups also emerged from the water cluster. One group was composed of microbial families from Actinomycetota, Pseudomonadota, Cyanobacteria, and other phyla. The second group was composed of families from ‘other’ microbial phyla, and interestingly, these microbial families were identified as significant in sawfish based on DA analysis. Therefore, these microbial families are contradictory because they exhibit a mean-abundance relationship that is higher in sawfish, but their abundance patterns correlate more closely with gene functions associated with the water column. The clustering of the sawfish families in the water column cluster suggests that these microbial families are ecological generalists occupying the sawfish ecological niche. The taxa–gene function correlation patterns suggest that the sawfish and the water column represent distinct ecosystems with unique ecological functions. This indicates that examining the relationships between taxa and their associated gene functions can provide predictable insights into the environmental characteristics shaping each community.

The uniform distribution of taxa and gene function correlation patterns within groups suggests potential functional redundancy. To examine this further, we identified the asymptotic behaviour of the taxa to gene function accumulation relationship. Across ten successively larger random sampling events of the dataset, we determined the taxa and gene function diversity accumulation (Figure 5). Over increasing sampling depths, the addition of new taxa resulted in faster gene function accumulation for the sawfish, relative to gene functions from the water column. Modeling this relationship with the Michaelis–Menten equation confirmed the trend, as indicated by the steeper rise of the sawfish curve (K = 201) compared with the water curve (K = 1622). In this context, K represents the half-saturation constant, defined as the number of taxa required to reach half of the maximum predicted gene functions, and thus quantifies how rapidly gene functions accumulate as taxa increases. The lower K value for the sawfish microbiome indicates a more efficient gain of functional diversity per taxon. Greater gene functions per taxa in the sawfish microbiome were further supported by the area under the curve (AUC), which agreed with the modeled results: sawfish gene functions accumulated more rapidly per taxa (Kruskal–Wallis: χ² = 9, df = 1, p < 0.005; AUC = 98,725 ± 1,170) compared with the water column (AUC = 84,246 ± 1,184), representing a 17.2% greater AUC for sawfish. Collectively, the Michaelis–Menten modeling and AUC analyses provide complementary evidence that the sawfish microbiome exhibits greater potential for functional redundancy. The lower K value reflects a more efficient accumulation of gene functions per taxon, while the higher AUC indicates a greater overall functional capacity, together supporting a functionally robust and overlapping microbial community.

**Figure 5:**
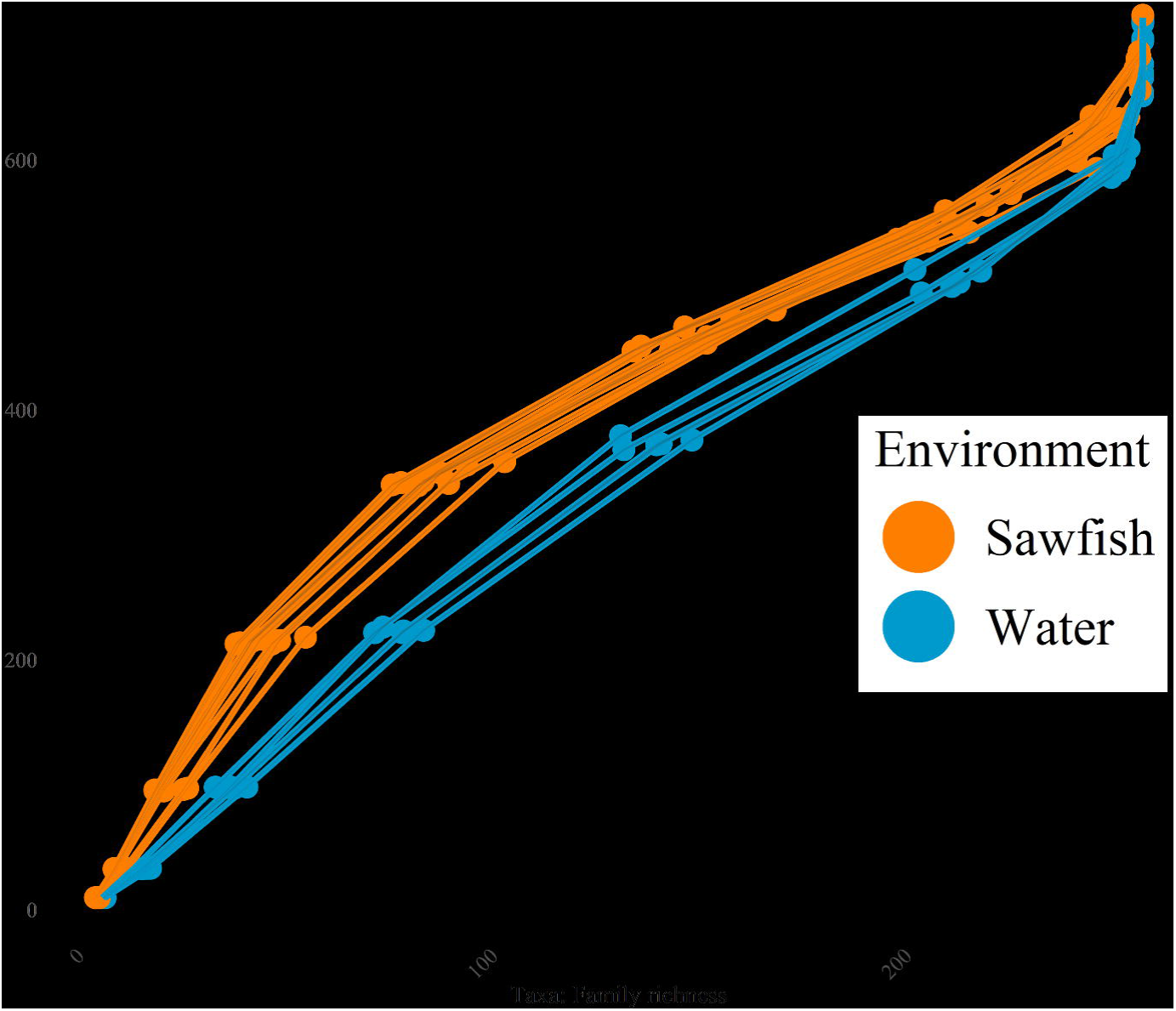
Accumulation of microbial family richness and gene function (Subsystem) richness with increasing sampling depth. Points represent the mean richness across 100 bootstrap resamples at each depth (library read depth) for significantly differentially abundant taxa and gene functions. Lines connect paired taxonomic and functional richness values within each sample and environment. Shaded ribbons represent ± standard error. At equivalent levels of functional richness, fewer microbial families are observed in the sawfish microbiome relative to the water column. For example, at approximately 200 gene functions, the sawfish microbiome contains fewer families than the water column, indicating greater taxonomic redundancy, where fewer taxa support a broader functional repertoire.

## 3. Discussion

The largetooth sawfish is a Critically Endangered euryhaline elasmobranch that in northern Australia can become isolated in drying floodplain pools in the late dry season. Extreme water quality in these evaporating pools can reduce their metabolic activity and increase susceptibility to environmental stressors and pathogen invasion. Here, we demonstrate that the sawfish skin microbiome is strongly host-associated and remains distinct from surrounding water column communities, despite individuals being sampled over multiple days. Dominant taxa and gene functions from these two environments demonstrate clear differences in microbiome composition. The water column microbiomes were more diverse compared with the sawfish. Enriched gene functions from the water include chlorophyll biosynthesis and gene corresponding to photosystems, suggesting energy production via photosynthesis. The taxonomic diversity of the sawfish skin microbiome is considerably lower than that of the surrounding water column, yet it harbors a distinct suite of gene functions with greater capacity for functional redundancy. Enrichment of genes associated with sporulation and anaerobic energy production suggests adaptation to the challenging conditions of the skin surface, where microbes likely utilize host-derived mucus as an energy source. Together, these results indicate that the sawfish skin supports a structured microbial community with unique metabolic capacities relative to the ambient environment. Integrating taxonomic and functional profiles provides clear evidence of host-driven filtering that selects for specific gene functions, supporting the hypothesis that the sawfish skin microbiome is functionally adapted to its host. These findings highlight the potential role of the microbiome in supporting host health and resilience under variable or stressful environmental conditions. Below, we discuss the ecology of the sawfish microbiome and its potential implications for sawfish health. We further discuss the role of microbiomes in the conservation of threatened species.

### 4.1 Lottery-like assembly of the sawfish microbiome

Understanding the ecological processes that shape host-associated microbiomes is critical for predicting how microbial communities influence host health, resilience, and conservation outcomes. For aquatic vertebrates, the mechanisms governing microbiome assembly remain largely unexplored, limiting our ability to anticipate responses to environmental change or to develop microbiome-informed conservation strategies. In this study, we show that the sawfish skin microbiome is highly host-specific, despite individuals being sampled over multiple days. Compared with the surrounding water column, sawfish support a lower richness of microbial families, suggesting strong host filtering, a more homogeneous set of niches on the host surface, or a combination of both. This pattern aligns with findings from freshwater fish in the Great Lakes (Sadeghi *et al*., 2023), although contrasting patterns have been observed in Amazonian species, where skin microbiomes were richer than the water column (Sylvain *et al*., 2020). Across each study, the skin microbiome is also consistently distinct from the surrounding environment, highlighting the role of both host and habitat in shaping community composition.

The sawfish microbiome also exhibits distinct functional characteristics: gene function profiles differ from the water column and individual taxa contribute a greater number of functions relative to environmental microbes, indicating high functional redundancy. Together, these patterns suggest that the sawfish microbiome assembles through a lottery-like mechanism. The Lottery Hypothesis posits that from a source pool, a subset of species with the functional capacity to occupy a given niche establishes stochastically, with priority effects governing which taxa persist, leading to high compositional variability despite functional similarity (Sale, 1976). Originally described for algal microbiomes (Burke *et al*., 2011) and later observed in thresher sharks (Doane *et al*., 2017), this mechanism contrasts with systems dominated by niche-driven assembly, such as the diatom *Thalassiosira rotula* (Mönnich *et al*., 2020). Functional redundancy in the sawfish microbiome, where many taxa share similar functions, supports the lottery model by linking taxonomic patterns to ecological roles. This redundancy illustrates how a host-specific microbiome can maintain functional stability while allowing stochastic assembly of taxonomically distinct taxa. Functional redundancy in this way can reduce the host’s reliance on its own immune system by enhancing colonization resistance of potential pathogens (Flores *et al*., 2025)

### 4.2 Bacillota and dormancy genes dominates the sawfish skin microbiome

The dominant microbial phylum of the sawfish microbiome was Bacillota (formerly Firmicutes) making up over 50% of the sawfish microbiome, while the water was comprised of a more diverse assemblage of evenly dominant microbial phyla, including Bacillota, Pseudomonadota, Actinomycota, and Bacteroidota. Bacillota is commonly found to be an abundant member of the gut microbiome (Ley *et al*., 2008) and in harsh environments (Filippidou *et al*., 2016). On the skin surface of aquatic animals, however, this bacterial phylum is not often found in high abundance. Typically, freshwater fish skin mucus microbiomes are dominated by Proteobacteria. For instance, Proteobacteria, on average, accounted for 78% of the skin microbiome spanning 17 freshwater fish species from the Great Lakes in the US (Sadeghi *et al*., 2023). Proteobacteria has been generally reported as the key elasmobranch microbiome phylum (Doane *et al*., 2020; Pogoreutz *et al*., 2025), while Bacillota remain in low abundance. For instance, in round stingrays (*Urobatis halleri*) and bat rays (*Myliobatis californica*), the microbiome was dominated by families from Proteobacteria and Bacteroidota rather than Bacillota (Kerr *et al*., 2023). This microbial phyla has been identified as discriminating microbiomes of fur seals from two locations, with increased Bacillota prevalence in locations with high seal density (Botsidou *et al*., 2025). Taxa from Bacillota were also described to be in high abundance in wound microbiomes of Indo-Pacific finless porpoise (*Neophocaena phocaenoides*) (Liu *et al*., 2022).

On sawfish, fifteen of the top twenty microbial families distinguishing the microbiome from the water column were families from Bacillota. The enriched Bacillota families in the sawfish microbiome are unique compared to families found in gut microbiomes, where Bacillota typically dominate. For instance, the gut microbiomes of farm animals and humans supported metagenome assembled genomes (MAGs) assigned to 37 Bacillota families, of which only five families from the sawfish overlapped (Machado et al., 2024). Bacillota are a phylum of Gram-positive bacteria often found in harsh conditions, a phenomenon attributed to their ability to form spores and persist through unfavorable conditions (Wörmer *et al*., 2019). However, sporulation only occurs within two classes—Bacilli and Clostridia (Galperin *et al*., 2022)—and the sawfish microbiome is dominated by taxa belonging to the class Bacilli.

The high abundance of Bacillota, and in particular families from the spore-forming class Bacilli suggest spore formation is a common phenomenon in the sawfish microbiome. Indeed, the microbiome is enriched for genes involved in sporulation and dormancy, suggesting that a significant portion of the microbial community can enter a quiescent state under environmental stress. The top five differentially abundant subsystems—spore germination, sporulation gene orphans, the SpoVA cluster, spore germinant receptors, and the SigE–G regulatory cluster—are canonical pathways for the formation, maintenance, and subsequent reactivation of bacterial endospores (Galperin *et al*., 2022). These functions are hallmarks of Bacillota and indicate that dormancy is a defining ecological strategy of this community. Their prevalence and dominance in the sawfish skin microbiome suggest that the skin surface environment imposes strong selective pressures consistent with a harsh, low-oxygen habitat. The enrichment of sporulation-related genes—including a two-fold increase in reads classified as spore germination and sporulation gene orphans, and a 1.4-fold enrichment of sporulation cluster genes—supports this interpretation. Spore formation provides clear advantages for resisting adverse physicochemical conditions and nutrient limitation (Wörmer *et al*., 2019). In other systems, such as the human gut, sporulation is well studied and hypothesized to be central to the persistence of Bacillota species (Browne *et al*., 2021). Moreover, sporulation may create a microbial reservoir that can repopulate once conditions improve, while maintaining community occupancy (i.e. niche filling) that limits pathogen invasion (Obeng *et al*., 2021).

Sporulation represents a critical adaptation for persistence in fluctuating or stressful environments. Its role in maintaining microbiome resilience is increasingly being recognized. For instance, a cultivation study using novel isolation methods revealed that 50–60% of human gut microbes harbor sporulation genes (Browne *et al*., 2016). For bacteria inhabiting the skin of sawfish from static, low-flow pools, oxygen diffusion is likely restricted, generating steep redox gradients and periodic energy limitation. Under such conditions, sporulation offers a mechanism for long-term survival when growth becomes energetically unfavorable. Spores are metabolically inert, resistant to desiccation and oxidative stress, and capable of germinating rapidly once favorable conditions return. The enrichment of spore-germination and regulatory gene clusters in the sawfish microbiome suggests not only the ability to endure dormancy but also a readiness to reinitiate metabolic activity, maintaining community continuity through variable oxygen availability and host activity cycles. This dormancy potential likely contributes to microbiome resilience during periods when the host’s own metabolic rate declines, such as during inactivity or stress. The ability of microbial populations to switch between active and dormant states could preserve community structure and functional potential even as nutrient and oxygen fluxes diminish.

Alongside the naturally low oxygen conditions of the pool, micro-layers forming at the skin interface can accentuate oxygen gradients, helping to explain the high abundance of Bacillota in the sawfish microbiome. These micro-layers create steep oxygen gradients (Roach *et al*., 2017), especially under low static flow conditions (Haas *et al*., 2014). The lack of flow, coupled with the heterotrophic activity of the microbes in these environments, mechanistically drives this phenomenon. The absence of shear stress exacerbates this gradient, limiting O□ exchange with the water column, and this can drastically alter biofilm communities in marine environments (Portas *et al*., 2024). Interestingly, Fang et al. (2017) showed that when soil microbial communities were exposed to different hydrostatic conditions—static versus flow (shear stress)—Bacillota dominated under static conditions, while Proteobacteria became dominant when flow was added. This suggests that shear forces can alter oxygen gradients that influence microbial community composition. By analogy, the skin surface of the sawfish may exhibit similar dynamics: relatively static flow conditions, coupled with microbial metabolic activity, likely create an oxygen-limited microenvironment. The dominance of Bacillota supports this prediction.

Adapting to the low oxygen environment in this way would ensure stability of the microbial community, which may be essential for host health. However, the present study was conducted on sawfish from a static floodplain pool system, where limited water movement likely amplifies oxygen gradients and favors sporulating, anaerobic taxa. Whether these traits are similarly advantageous in free-flowing, oxygen-rich environments remains an open question. Investigating individuals from river or estuarine environments will clarify whether Bacillota dominance and dormancy-associated functions are intrinsic to the sawfish microbiome or represent a context-dependent response to low flow conditions. Understanding these dynamics is crucial, as resilient microbial communities could play an unrecognized role in supporting the health, environmental tolerance, and conservation of Critically Endangered species like the largetooth sawfish.

### 4.3 Anaerobic metabolism of sawfish mucus by the microbiome

Consistent with the ecological interpretation of an oxygen-limited skin environment, the functional gene repertoire of the microbiome indicates that energy generation is largely sustained through the metabolism of host-derived mucus substrates. From enriched gene functions, two distinct pathways for energy generation converge on a central energy source: host mucus. Mucus is composed mainly of mucins, which are glycoproteins with a protein backbone densely decorated with glycan chains derived from monosaccharides (Bansil and Turner, 2018), of which each can provide a carbon rich media with large energy potential to support microbial metabolism (Wu *et al*., 2023). Genes associated with the putative metabolism of mucus glycans were enriched in the skin microbiomes of southern eagle rays (*Myliobatis tenuicaudatus*) relative to water column microbial communities (Kerr *et al*., 2026). When metabolized, glycoproteins are broken down into dipeptides (Winchester, 2005), and dipeptide ABC transporters are enriched in the sawfish microbiome, indicating the potential for uptake by skin microbes of these substrates. Imported dipeptides are then hydrolyzed into amino acids, which can be transaminated into α-keto acids (i.e. pyruvate). Gene functions for the utilization of branched-chain amino acids and α-keto acids are enriched in this environment, suggesting the high potential for this process to be occurring. These α-keto acids are then oxidatively decarboxylated to form acyl-CoA derivatives, which can enter either the TCA cycle or acetogenesis/fermentation pathways for energy production. The enrichment of pyruvate formate-lyase in the sawfish microbiome supports the anaerobic pyruvate metabolism to acetyl-CoA for acetogenesis (Krivoruchko *et al*., 2015).

In addition, uronic acid utilization pathways, specifically for the metabolisms of D-Galacturonate and D-Glucuronate, are enriched, suggesting mucus glycans are also metabolized by the sawfish microbiome (Yamaguchi and Yamamoto, 2023). Uronic acids can be readily converted to acetyl-CoA via pyruvate for potential acetogenesis (Hager *et al*., 2019). While gene functions involved in the TCA cycles are also abundant in the sawfish microbiome, acetogenesis-related functions are substantially more abundant in the sawfish microbiome. This strongly supports anaerobic acetogenesis as the dominant energy-generating pathway in the sawfish microbiome. These results further support the theory that the skin microbiome inhabits an O2-depleted micro-habitat, described above. A key unresolved question is whether the observed microbiome represents an adapted microbiome in a low oxygen environment, or whether the microbiome constitutes the normal adapted community despite the low oxygen concentration in the pool. For this, we require samples to be taken from individuals occupying free-flowing environments with elevated oxygen concentrations. Despite this, the microbiome is indeed selective and adapted to similar conditions, as we describe microbiome patterns from individuals over time points with varying oxygen saturation, with the microbiome converging. In humans, mucus glycan can provide disease insights, acting as biomarkers (Modesto *et al*., 2023). Our results demonstrate close links between the sawfish mucus and its microbiome, which stands to reason that the microbiome will be a bioindicator of sawfish physiological status.

### 4.4 The sawfish microbiome in conservation

For species of concern, enhancing fitness is paramount for interventions aimed at preventing extinction. Microbial communities living with host animals substantially influence host fitness, both directly through metabolite production and indirectly by shaping host phenotype, that enables adaptation and resilience to environmental change (Morais *et al*., 2021). In aquatic environments, a host’s microbiome maintains a distinct composition from the surrounding water column microbial communities that can closely mirror environmental conditions (e.g., Amazonian fish species; (Sehnal *et al*., 2021)) or be largely species-specific (e.g., coral reef fishes; (Chiarello *et al*., 2015)), reflecting both ecological and evolutionary constraints. In this way, the microbiome acts as an extension of the immune system, which can minimize colonization by potentially harmful taxa (Spragge *et al*., 2023).

Microbiomes can contribute to physiological plasticity in wild fish with low genetic diversity (Anka *et al*., 2024), a critical factor for species like sawfish with small, fragmented populations. Recognizing the microbiome’s adaptive role and its health consequences for the host, Mueller et al. (2020) proposed a framework for incorporating microbiomes into conservation strategies — an approach especially critical for threatened species. According to the microbial rescue framework, beneficial microbial associations can promote host persistence under environmental change through three stages: (1) environmentally induced shifts in microbiome composition or function, (2) host fitness benefits through conferred microbiome attributes (e.g., microbial products that regulate inflammation via host immune cells), and (3) transmission of advantageous microbial traits among individuals, ultimately supporting population recovery.

Developing the foundation for how the microbiome assembles and adapts underpins the microbiome’s role in host fitness. Our findings present baseline insight into the skin microbiome dynamics and demonstrate a distinct, host-associated microbiome community capable of maintaining metabolic activity under low-oxygen and nutrient-limited conditions. Sawfish experience dynamic environmental conditions that span varying degrees of nutrient and oxygen availability to its microbiome. The dominance of Bacillota and enrichment of sporulation and anaerobic energy pathways suggest a microbiome well adapted to persist through these expected prolonged environmental stresses. Derived from these results, we hypothesize the microbiome is an active extension of the sawfish’s physiological barrier to the external environment. This resilience implies that the microbiome is not merely a passive reflection of environmental conditions, but an active extension of the host’s physiology, contributing significantly to the host’s immune system and microbiome stability.

Maintaining stability and stress-tolerance enables the microbiome to be a biological barrier that can limit colonization by opportunistic microbes. Consequently, the stability and stress tolerance of the microbiome provide a unique window into the host’s biochemical responses to stress. For instance, if mucus production or the composition of glycan decorating the glycoproteins shifts in response to the host’s physiological stress, corresponding changes in the microbiome could serve as early, non-invasive indicators of environmental disturbance or declining health. As a direct source of energy, mucus is a key link between the microbiome and the physiological state of the host. Microbiome compositional shifts occur as a direct response to distinct glycosylation patterns (Wu et al., 2023). Integrating microbiome monitoring into conservation frameworks is, therefore, warranted to more effectively enhance our ability to detect and mitigate stress responses to species of concern before they manifest at the population level.

## 4. Conclusion

Together, these results reveal that the sawfish skin supports a microaerobic-to-anaerobic environment that selects for a metabolically specialized microbiome. The dominance of Bacillota, coupled with enrichment of gene functions for sporulation, anaerobic energy metabolism, and mucus degradation, indicates that microbial life on the sawfish skin is sustained by host-derived substrates under low-oxygen conditions. The patterns of functional redundancy observed in this microbiome suggest a lottery-like mechanism of community assembly, where multiple, taxonomically distinct taxa can stochastically occupy ecological niches while maintaining key functional capabilities. This mechanism likely supports microbiome stability during periods of environmental stress, a characteristic that can mediate pathogen invasion and enhance host resilience. These findings strongly indicate that the sawfish microbiome has potential as an early indicator of host health for monitoring purposes. Importantly, these observations are based on sawfish individuals in a static, pool environment, where limited water flow likely amplifies oxygen gradients and shapes microbial composition. Future work should examine sawfish from free-flowing habitats to determine whether Bacillota dominance persists or if community assembly shifts toward other taxa under more oxygenated conditions. Understanding these dynamics is crucial, as microbiomes that enhance host resilience to environmental stressors may play an underappreciated role in the health and conservation of Critically Endangered species like the largetooth sawfish.

## Ethics Statement

Activities were performed under Flinders University ethics AEC BIOL9436-2.

## Data Availability

All scripts used for analysis and figure generation are available at https://github.com/mpdoane2/Sawfish-microbiome-analysis. Raw metagenomic sequence data are available through the NCBI Sequence Read Archive (SRA) under BioProject accession number PRJNA1431019.

## Acknowledgements

We thank Karen Gibb, Keith Christian, and Zarah Tinning (Research Institute for the Environment and Livelihoods, Charles Darwin University) for project and sampling assistance. Funding was provided by the Charles Darwin University 2021 Institute of Advanced Studies (IAS) Rainmaker Start Up and the Australian Marine Conservation Society (AMCS). We thank Tooni Mahto of AMCS for project support. We further thank the Flinders Accelerator for Microbiome Exploration (FAME) for financially supporting the metagenomic library generation and sequencing efforts.

## Author Contributions

**Michael P. Doane:** Conceptualization (lead), data curation (equal), formal analysis (lead), investigation(lead), methodology(supporting), visualization(lead), writing – original draft preparation(lead)/Review and editing(equal). **Belinda Martin:** methodology (lead), writing – original draft (supporting)/review and editing (equal). **Emma Kerr:** methodology (supporting), writing – original draft(supporting)/review and editing (equal). **Elizabeth A. Dinsdale:** Investigation (supporting), supervision (supporting), writing – original draft (supporting)/review and editing (equal). **Malak Malak Rangers:** data curation (equal), writing – review and editing (equal). **Leonardo Guida**: data curation (equal), writing – review and editing (equal). **Pete M. Kyne:** Conceptualization (supporting), data curation (equal), writing – original draft preparation (supporting)/review and editing (equal).

**SI Figure 1:** Proportion of host-filtered metagenomic reads assigned to a taxonomy and a SEED Subsystem gene function for largetooth sawfish (*Pristis pristis*) skin and water samples from the Daly River floodplain, Northern Territory, Australia. (A) represents the taxonomic assignments and (B) represents the gene function assignments. The upper and lower bounds of the box represent the first and third quantiles, with the median value in the box. The whiskers note the minimum and maximum values with outliers displayed as dots beyond the whiskers. The dots within each plot correspond to the sample value to show data distribution.

**SI Figure 2:** Shared taxonomic groups from the sawfish and the water column microbiome. The total number of shared groups (Gamma diversity richness) is on top with percent in parentheses. Percentage is based on the total (Gamma diversity) across the combined dataset. The total count of taxa unique to sawfish is in blue, and the water column is in yellow. The share portion is the values in the overlapping section. (A) represents the shared taxonomy at the family level, while (B) represents the genus level.

**SI Figure 3:** The distribution of microbiome richness from individual samples across the two environments (sawfish; water). Richness is measured as the number of unique microbial species identified in the dataset. Circles represent samples sequenced using 2×150 bp inserts, while triangles represent 2×300 bp sequencing. The red dashed line corresponds to the mean richness across all samples (mean alpha diversity), and the blue dashed line corresponds to the gamma diversity, defined as the total number of unique species across each dataset. The size of the shape is representative of the library size.

**SI Figure 4:** Boxplots showing family-level Shannon diversity of microbial communities associated with sawfish skin and the surrounding water column. Boxes represent the interquartile range with the median indicated by the central line; whiskers extend to 1.5 × the interquartile range, and points denote individual samples. Differences between host-associated and free-living communities highlight contrasts in diversity structure between the sawfish skin microbiome and the ambient water column.

**SI Figure 5:** Microbial distribution of the top 20 microbial Families in sawfish and surrounding water column microbial communities at the taxonomic level of **f**amily. The x-axis are the samples from the sawfish and water. The y-axis represents the proportion of the total number of sequencing reads. The figure legend indicates the level which the microbial group is, for instance k indicates Kingdom, p Phylum, o Order, and f Family. Read sequences that were mapped equally across Families were assigned the common Order level, or Phylum. All other microbial Families are assigned to Low abundant.

**SI Figure 6:** The microbial family level of the sawfish and water microbiomes represented by nMDS ordination plot. The distance between any two samples corresponds to the relative similarity in microbiome composition, measured by Bray-Curtis similarity from 4th-root transformed relative abundance data. Colour represents the day of sampling and the shape corresponds to the sequencing library size.

**SI Figure 7:** Richness of Subsystems across Level 1 SEED functional categories. Richness represents the number of Subsystems detected within each Level 1 category for sawfish and water column environments. Error bars indicate standard error. The x-axis shows the number of Subsystems, and the y-axis lists the Level 1 functional categories.

**SI Figure 8:** Abundance of Subsystems across Level 1 SEED functional categories. Abundance is represented as the number of sequencing reads assigned to each Subsystem within a Level 1 category for sawfish and water column environments. Error bars indicate standard error. The x-axis shows read count (abundance), and the y-axis lists the Level 1 functional categories.

**SI Figure 9:** The gene functions of the sawfish and water microbiomes represented by nMDS ordination plot. The functions in sawfish and water microbiomes were distinctive, with high similarity among most replicate samples (samples from similar environments). Color represents the day and environment from which the sample was taken. Shape corresponds to the read length. The distance between any two samples corresponds to the relative similarity in microbiome composition, measured by Bray-Curtis similarity from 4th-root transformed relative abundance data.

**SI Figure 10:** nMDS ordination of microbial energy subsystems from sawfish and water column environments. Circles and triangles represent metagenomic libraries of different sequence lengths, illustrating that environment is the primary driver of microbiome gene function patterns. Vectors indicate the direction of positive influence of each subsystem and are colored by their corresponding Level 3 category. Stress values indicate the goodness-of-fit of the nMDS ordination.

**SI Figure 11:** Heatmap showing the abundance relationship between all microbial taxa and gene functions from the sawfish and water column microbiomes. Red and blue correspond to positive and negative Spearman correlation, respectively. Lighter colors signify a weak relationship. On the x-axis, taxa are labelled by the phyla in which they are classified and the group from which they are significantly enriched. Gene functions are represented by rows and organized by their mean abundance across the dataset within the Level 1 SEED category (the broadest classification).

**SI Table 1:** Sample metadata for sawfish skin and water-column microbiomes. Metadata include sample ID, collection site, date, environmental parameters (e.g., O□ saturation, temperature, flow), and sample type (skin or water column).

**SI Table 2:** ANCOM-BC results for differential abundance of microbial taxa at the Family level. This table reports all Families detected in sawfish skin and water-column microbiomes, including ANCOM-BC log fold changes, standard errors, test statistics, p-values, and adjusted p-values for multiple testing. Positive LFC values correspond to the water, and negative LFC is the sawfish environment.

**SI Table 3:** ANCOM-BC results for differential abundance of microbial gene functions at the Subsystem level. This table lists all functional gene Subsystems detected in sawfish skin and water-column metagenomes, including ANCOM-BC log fold changes, standard errors, test statistics, p-values, and adjusted p-values for multiple testing. Positive LFC values correspond to the water, and negative LFC is the sawfish environment.

---

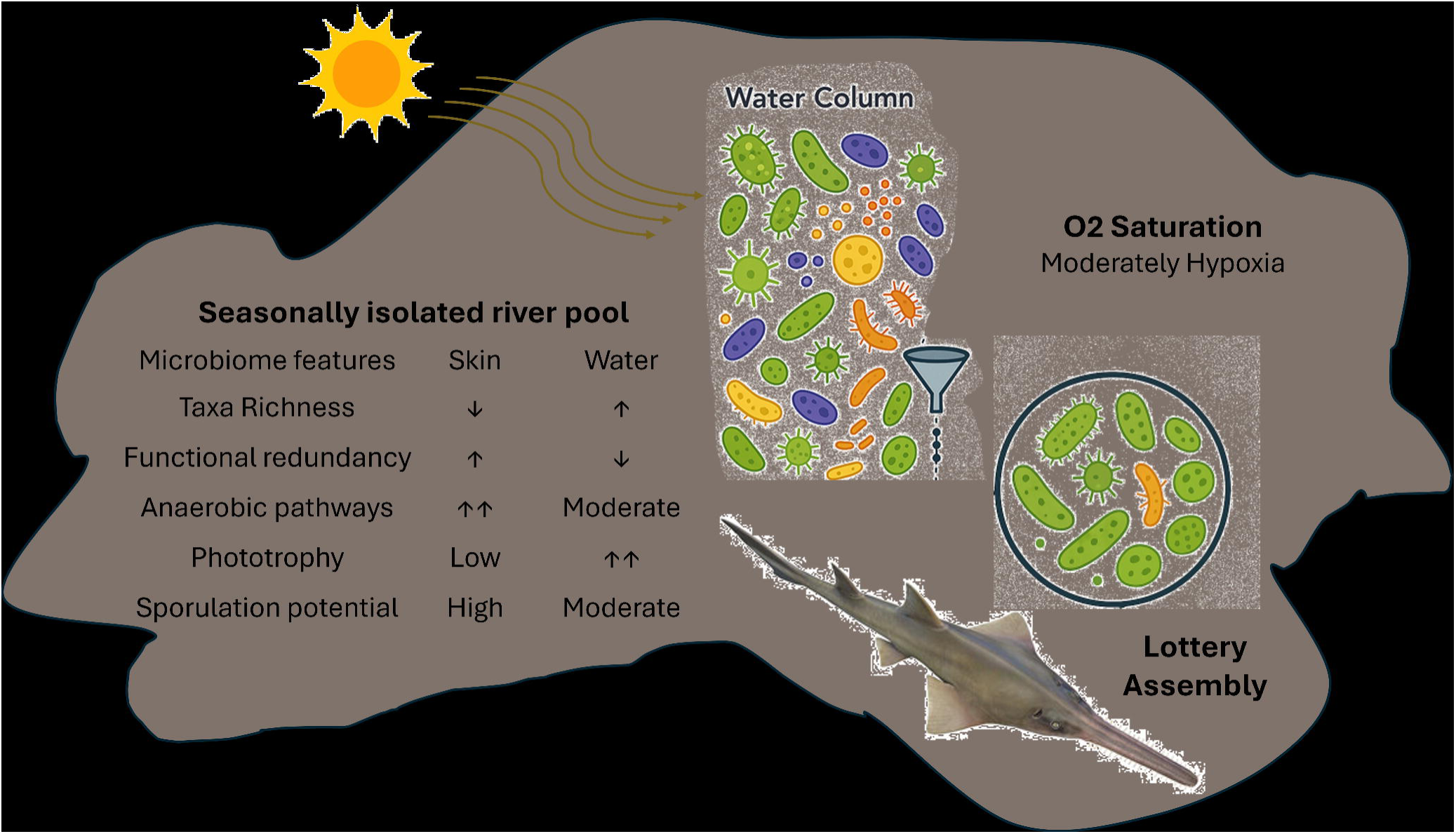

## Bibliographies

Albert, J.S., Destouni, G., Duke-Sylvester, S.M., Magurran, A.E., Oberdorff, T., Reis, R.E., et al. (2021) Scientists’ warning to humanity on the freshwater biodiversity crisis. Ambio 50: 85–94.

Anderson, M.J. (2001) A new method for non parametric multivariate analysis of variance. Austral Ecol 26: 32–46.

Anka, I.Z., Uren Webster, T.M., Berbel-Filho, W.M., Hitchings, M., Overland, B., Weller, S., et al. (2024) Microbiome and epigenetic variation in wild fish with low genetic diversity. Nat Commun 15: 4725.

Apprill, A., Miller, C.A., van Cise, A.M., U’Ren, J.M., Leslie, M.S., Weber, L., et al. (2020) Marine mammal skin microbiotas are influenced by host phylogeny. R Soc Open Sci 7: 192046.

Bansil, R. and Turner, B.S. (2018) The biology of mucus: Composition, synthesis and organization. Adv Drug Deliv Rev 124: 3–15.

Bell, A.G., McMurtrie, J., Bolaños, L.M., Cable, J., Temperton, B., and Tyler, C.R. (2024) Influence of host phylogeny and water physicochemistry on microbial assemblages of the fish skin microbiome. FEMS Microbiol Ecol 100: fiae021.

Borchers, H.W. (2025) pracma: Practical Numerical Math Functions. R Packag version 246.

Bordenstein, S.R., The Holobiont Biology Network, Gilbert, M.T.P., Ginnan, N., Malacrinò, A., Martino, M.E., et al. (2024) The disciplinary matrix of holobiont biology. Science (80-) 386: 731–732.

Botsidou, P., Schloter, M., Maraci, Ö., Gschwendtner, S., Nagel, R., Forcada, J., and Hoffman, J.I. (2025) Skin, but not gut, microbial communities vary with social density in Antarctic fur seals. Front Microbiol 16: 1603500.

Brown, A.L., Sharp, K., and Apprill, A. (2022) Reshuffling of the Coral Microbiome during Dormancy. Appl Environ Microbiol 88: 01291–22.

Browne, H.P., Almeida, A., Kumar, N., Vervier, K., Adoum, A.T., Viciani, E., et al. (2021) Host adaptation in gut Firmicutes is associated with sporulation loss and altered transmission cycle. Genome Biol 22: 1–20.

Browne, H.P., Forster, S.C., Anonye, B.O., Kumar, N., Neville, B.A., Stares, M.D., et al. (2016) Culturing of ‘unculturable’ human microbiota reveals novel taxa and extensive sporulation. Nature 533: 1–15.

Burke, C., Steinberg, P., Rusch, D., Kjelleberg, S., and Thomas, T. (2011) Bacterial community assembly based on functional genes rather than species. PNAS 108: 14288–14293.

Cavicchioli, R., Ripple, W.J., Timmis, K.N., Azam, F., Bakken, L.R., Baylis, M., et al. (2019) Scientists’ warning to humanity: microorganisms and climate change. Nat Rev Microbiol 17: 569–586.

Chiarello, M., Auguet, J.-C., Bettarel, Y., Bouvier, C., Claverie, T., Graham, N.A.J., et al. (2018) Skin microbiome of coral reef fish is highly variable and driven by host phylogeny and diet. Microbiome 6: 147.

Chiarello, M., Villéger, S., Bouvier, C., Bettarel, Y., and Bouvier, T. (2015) High diversity of skin-associated bacterial communities of marine fishes is promoted by their high variability among body parts, individuals and species. FEMS Microbiol Ecol 91: fiv061.

Darwall, W., Bremerich, V., De Wever, A., Dell, A.I., Freyhof, J., Gessner, M.O., et al. (2018) The Alliance for Freshwater Life: A global call to unite efforts for freshwater biodiversity science and conservation. Aquat Conserv Mar Freshw Ecosyst 28: 1015–1022.

Dinno, A. (2014) dunn.test: Dunn’s Test of Multiple Comparisons Using Rank Sums. CRAN Contrib Packag.

Dinsdale, E.A., Edwards, R.A., Hall, D., Angly, F., Breitbart, M., Brulc, J.M., et al. (2008) Functional metagenomic profiling of nine biomes. Nature 452: 629–32.

Doane, M.P., Haggerty, J.M., Kacev, D., Papudeshi, B., and Dinsdale, E.A. (2017) The skin microbiome of the common thresher shark (*Alopias vulpinus*) has low taxonomic and gene function β[diversity. Environ Microbiol Rep 9: 357–373.

Doane, M.P., Johnson, C.J., Johri, S., Kerr, E.N., Morris, M.M., Desantiago, R., et al. (2022) The Epidermal Microbiome Within an Aggregation of Leopard Sharks (*Triakis semifasciata*) Has Taxonomic Flexibility with Gene Functional Stability Across Three Time-points. Microb Ecol 85: 747–764.

Doane, M.P., Morris, M.M., Papudeshi, B., Allen, L., Pande, D., Haggerty, J.M., et al. (2020) The skin microbiome of elasmobranchs follows phylosymbosis, but in teleost fishes the microbiomes converge. Microbiome 8: 1–15.

Doane, M.P., Ostrowski, M., Brown, M., Bramucci, A., Bodrossy, L., van de Kamp, J., et al. (2023) Defining marine bacterioplankton community assembly rules by contrasting the importance of environmental determinants and biotic interactions. Environ Microbiol 25: 1084–1098.

Doane, M.P., Reed, M.B., McKerral, J., Farias Oliveira Lima, L., Morris, M., Goodman, A.Z., et al. (2023) Emergent community architecture despite distinct diversity in the global whale shark (*Rhincodon typus*) epidermal microbiome. Sci Rep 13: 12747.

Edwards, R.A. (2025) linsalrob/atavide_lite: Ecology in Focus Release (version v0.3). Zenodo.

Ernst, F.G.M., Borman, S.A.S.T., and Lahti, L. (2023) mia: Microbiome analysis.

Espinoza, M., Bonfil-Sanders, R., Carlson, J., Charvet, P., Chevis, M., Dulvy, N.K., et al. (2022) Pristis pristis. IUCN Red List Threat Species e-T18584848A.

Filippidou, S., Wunderlin, T., Junier, T., Jeanneret, N., Dorador, C., Molina, V., et al. (2016) A Combination of Extreme Environmental Conditions Favor the Prevalence of Endospore-Forming Firmicutes. Front Microbiol 7: 1707.

Flinders University High-Performance Computing cluster (2021) DeepThought (HPC).

Flores, C., Millard, S., and Seekatz, A.M. (2025) Bridging Ecology and Microbiomes: Applying Ecological Theories in Host-associated Microbial Ecosystems. Curr Clin Microbiol Reports 12: 9.

Galperin, M.Y., Yutin, N., Wolf, Y.I., Alvarez, R.V., and Koonin, E. V. (2022) Conservation and Evolution of the Sporulation Gene Set in Diverse Members of the Firmicutes. J Bacteriol 204: e00079–22.

Goodman, A.Z., Papudeshi, B., Doane, M.P., Mora, M., Kerr, E., Torres, M., et al. (2022) Epidermal microbiomes of leopard sharks (*Triakis semifasciata*) are consistent across Captive and wild environments. Microorganisms 10: 2081.

Haas, A.F., Smith, J.E., Thompson, M., and Deheyn, D.D. (2014) Effects of reduced dissolved oxygen concentrations on physiology and fluorescence of hermatypic corals and benthic algae. PeerJ 2: e235.

Hager, F.F., Sützl, L., Stefanović, C., Blaukopf, M., and Schäffer, C. (2019) Pyruvate Substitutions on Glycoconjugates. Int J Mol Sci 20: 4929.

IUCN (2025) The IUCN Red List of Threatened Species. IUCN Red List Threat Species Version 2025-2 https://www.iucnredlist.org Accessed on 10 July 2025.

Keck, F., Peller, T., Alther, R., Barouillet, C., Blackman, R., Capo, E., et al. (2025) The global human impact on biodiversity. Nature 641: 395–400.

Kerr, E.N., Papudeshi, B., Haggerty, M., Wild, N., Goodman, A.Z., Lima, L.F.O., et al. (2023) Stingray epidermal microbiomes are species-specific with local adaptations. Front Microbiol 14: 1–13.

Kerr, E.N., Yu, L., Hesse, R.D., Roberts, C.N., Bulone, V., Meyer, L., et al. (2026) Interactions of mucus monosaccharides and the epidermal microbiome in four benthic Elasmobranchs. Environ Microbiol Rep Advance online publication. 10.1111/1758-2229.70303.

Kolde, R. (2025) pheatmap: Pretty Heatmaps. R Packag version 1013.

Krivoruchko, A., Zhang, Y., Siewers, V., Chen, Y., and Nielsen, J. (2015) Microbial acetyl-CoA metabolism and metabolic engineering. Metab Eng 28: 28–42.

Lear, K.O., Morgan, D.L., Whitty, J.M., Beatty, S.J., and Gleiss, A.C. (2021) Wet season flood magnitude drives resilience to dry season drought of a euryhaline elasmobranch in a dry-land river. Sci Total Environ 750: 142234.

Ley, R.E., Hamady, M., Lozupone, C., Turnbaugh, P.J., Ramey, R.R., Bircher, J.S., et al. (2008) Evolution of Mammals and Their Gut Microbes. Science 320: 1647–1651.

Lin, H. and Peddada, S. Das (2020) Analysis of compositions of microbiomes with bias correction. Nat Commun 11: 3514.

Liu, Z., Wang, J., Meng, D., Li, L., Liu, X., Gu, Y., et al. (2022) The self-organization of marine microbial networks under evolutionary and ecological processes: Observations and modeling. Biology (Basel) 11: 592.

Louca, S., Parfrey, L.W., and Doebeli, M. (2016) Decoupling function and taxonomy in the global ocean microbiome. Science 353: 1272–1277.

McCauley, D.J., Pinsky, M.L., Palumbi, S.R., Estes, J.A., Joyce, F.H., and Warner, R.R. (2015) Marine defaunation: Animal loss in the global ocean. Science 347: 1255641–1.

McMurdie, P.J. and Holmes, S. (2013) phyloseq: An R Package for Reproducible Interactive Analysis and Graphics of Microbiome Census Data. PLoS One 8: e61217.

Modesto, J.L., Pearce, V.H., and Townsend, G.E. (2023) Harnessing gut microbes for glycan detection and quantification. Nat Commun 14: 275.

Mönnich, J., Tebben, J., Bergemann, J., Case, R., Wohlrab, S., and Harder, T. (2020) Niche-based assembly of bacterial consortia on the diatom *Thalassiosira rotula* is stable and reproducible. ISME J 14: 1614–1625.

Morais, L.H., IV, H.L.S., and Mazmanian, S.K. (2021) The gut microbiota – brain axis in behaviour and brain disorders. Nat Rev Microbiol 19: 241–255.

Obeng, N., Bansept, F., Sieber, M., Traulsen, A., and Schulenburg, H. (2021) Evolution of Microbiota–Host Associations: The Microbe’s Perspective. Trends Microbiol 29: 779–787.

Oksanen, J., Blanchet, F.G., Friendly, M., Kindt, R., Legendre, P., McGlinn, D., et al. (2020) vegan: Community Ecology Package. R Packag version 25-7.

Olson, R.D., Assaf, R., Brettin, T., Conrad, N., Cucinell, C., Davis, J.J., et al. (2023) Introducing the Bacterial and Viral Bioinformatics Resource Center (BV-BRC): a resource combining PATRIC, IRD and ViPR. Nucleic Acids Res 51: D678–D689.

Opstal, E.J. Van and Bordenstein, S.R. (2015) Rethinking heritability of the microbiome. Science 349: 1172–1173.

Peixoto, R.S., Voolstra, C.R., Sweet, M., Duarte, C.M., Carvalho, S., Villela, H., et al. (2022) Harnessing the microbiome to prevent global biodiversity loss. Nat Microbiol 7: 1726–1735.

Pogoreutz, C., Gore, M., Perna, G., Ormond, R., Clarke, C.R., and Voolstra, C.R. (2025) Microenvironments of black-tip reef sharks (*Carcharhinus melanopterus*) provide niche habitats for distinct bacterial communities. Coral Reefs 44: 145–162.

Portas, A., Carriot, N., Barry-Martinet, R., Ortalo-Magné, A., Hajjoul, H., Dormoy, B., et al. (2024) Shear stress controls prokaryotic and eukaryotic biofilm communities together with EPS and metabolomic expression in a semi-controlled coastal environment in the NW Mediterranean Sea. Environ Microbiome 19: 109.

Regan, M.D., Chiang, E., Liu, Y., Tonelli, M., Verdoorn, K.M., Gugel, S.R., et al. (2022) Nitrogen recycling via gut symbionts increases in ground squirrels over the hibernation season. Science 375: 460–463.

Reid, A.J., Carlson, A.K., Creed, I.F., Eliason, E.J., Gell, P.A., Johnson, P.T.J., et al. (2019) Emerging threats and persistent conservation challenges for freshwater biodiversity. Biol Rev 94: 849–873.

Roach, T.N.F., Abieri, M.L., George, E.E., Knowles, B., Naliboff, D.S., Smurthwaite, C.A., et al. (2017) Microbial bioenergetics of coral-algal interactions. PeerJ 2017: 1–19.

Ross, E.R. and Randhir, T.O. (2022) Effects of climate and land use changes on water quantity and quality of coastal watersheds of Narragansett Bay. Sci Total Environ 807: 151082.

Ruiz-Rodríguez, M., Scheifler, M., Sanchez-Brosseau, S., Magnanou, E., West, N., Suzuki, M., et al. (2020) Host Species and Body Site Explain the Variation in the Microbiota Associated to Wild Sympatric Mediterranean Teleost Fishes. Microb Ecol 80: 212–222.

Sadeghi, J., Chaganti, S.R., Johnson, T.B., and Heath, D.D. (2023) Host species and habitat shape fish-associated bacterial communities: phylosymbiosis between fish and their microbiome. Microbiome 11: 1–19.

Sale, P.F. (1976) Reef fish lottery. Nat Hist 85: 60–65.

Sehnal, L., Brammer-Robbins, E., Wormington, A.M., Blaha, L., Bisesi, J., Larkin, I., et al. (2021) Microbiome Composition and Function in Aquatic Vertebrates: Small Organisms Making Big Impacts on Aquatic Animal Health. Front Microbiol 12: 567408.

Shen, W. and Ren, H. (2021) TaxonKit: A practical and efficient NCBI taxonomy toolkit. J Genet Genomics 48: 844–850.

Spragge, F., Bakkeren, E., Jahn, M.T., Araujo, E.B.N., Pearson, C.F., Wang, X., et al. (2023) Microbiome diversity protects against pathogens by nutrient blocking. Science 382: eadj3502.

Sylvain, F.-É., Holland, A., Bouslama, S., Audet-Gilbert, É., Lavoie, C., Val, A.L., and Derome, N. (2020) Fish Skin and Gut Microbiomes Show Contrasting Signatures of Host Species and Habitat. Appl Environ Microbiol 86: e00789–20.

Troitsky, T.S., Laine, V.N., and Lilley, T.M. (2023) When the host’s away, the pathogen will play: the protective role of the skin microbiome during hibernation. Anim Microbiome 5: 66.

Turnbaugh, P.J., Ley, R.E., Hamady, M., Fraser-Liggett, C.M., Knight, R., and Gordon, J.I. (2007) The human microbiome project. Nature 449: 804–810.

Walling, L.K., Gamache, M.H., González-Pech, R.A., Harwood, V.J., Ibrahim-Hashim, A., Jung, J.H., et al. (2025) Incorporating microbiome analyses can enhance conservation of threatened species and ecosystem functions. Sci Total Environ 970: 178826.

Wickham, H. (2016) ggplot2: Elegant Graphics for Data Analysis. Springer-Verlag New York.

Wilde, J., Slack, E., and Foster, K.R. (2024) Host control of the microbiome: Mechanisms, evolution, and disease. Science (80-) 385: eadi3338.

Winchester, B. (2005) Lysosomal metabolism of glycoproteins. Glycobiology 15: 1–15.

Wörmer, L., Hoshino, T., Bowles, M.W., Viehweger, B., Adhikari, R.R., Xiao, N., et al. (2019) Microbial dormancy in the marine subsurface: Global endospore abundance and response to burial. Sci Adv 5: 24–26.

Wu, C.M., Wheeler, K.M., Cárcamo-Oyarce, G., Aoki, K., McShane, A., Datta, S.S., et al. (2023) Mucin glycans drive oral microbial community composition and function. npj Biofilms Microbiomes 9: 11.

Wu, G., Xu, T., Zhao, N., Lam, Y.Y., Ding, X., Wei, D., et al. (2024) A core microbiome signature as an indicator of health. Cell 187: 6550–6565.e11.

Yamaguchi, M. and Yamamoto, K. (2023) Mucin glycans and their degradation by gut microbiota. Glycoconj J 40: 493–512.

