## Supplemental Figures for "Functional redundancy and anaerobic metabolism characterize the skin microbiome of a critically endangered sawfish"

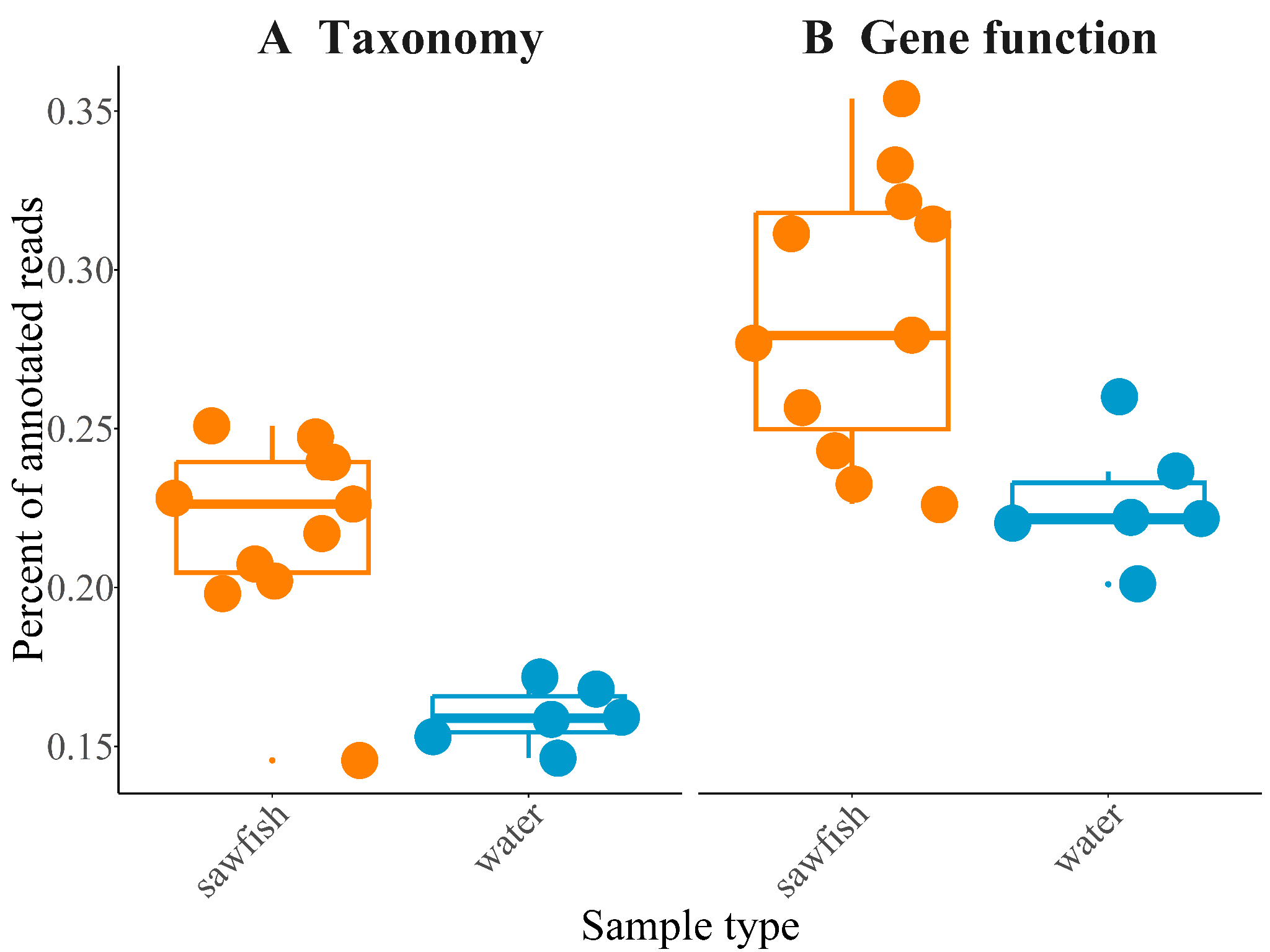

**SI Figure 1:** Proportion of host-filtered metagenomic reads assigned to a taxonomy and a SEED Subsystem gene function for largetooth sawfish (*Pristis pristis*) skin and water samples from the Daly River floodplain, Northern Territory, Australia. (A) represents the taxonomic assignments and (B) represents the gene function assignments. The upper and lower bounds of the box represent the first and third quantiles, with the median value in the box. The whiskers note the minimum and maximum values with outliers displayed as dots beyond the whiskers. The dots within each plot correspond to the sample value to show data distribution.

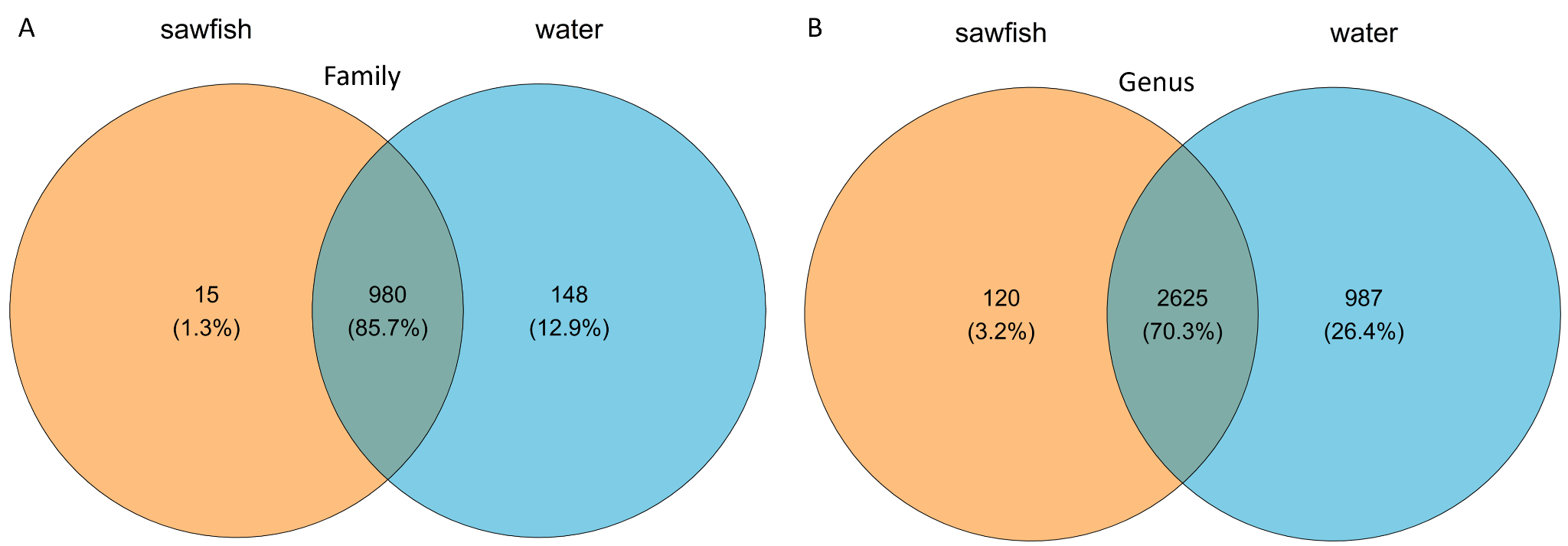

**SI Figure 2:** Shared taxonomic groups from the sawfish and the water column microbiome. The total number of shared groups (Gamma diversity richness) is on top with percent in parentheses. Percentage is based on the total (Gamma diversity) across the combined dataset. The total count of taxa unique to sawfish is in blue, and the water column is in yellow. The share portion is the values in the overlapping section. (A) represents the shared taxonomy at the family level, while (B) represents the genus level.
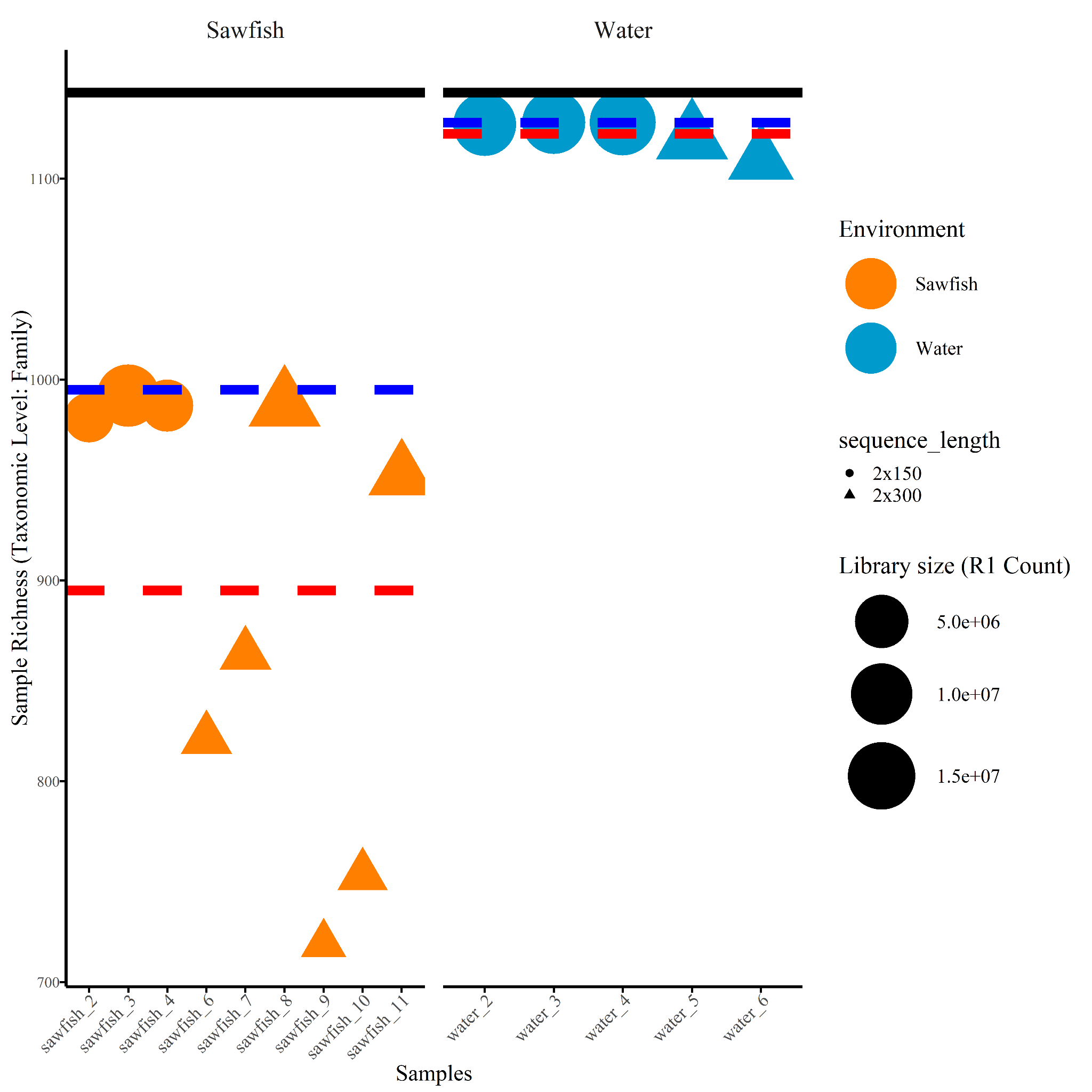

**SI Figure 3***:* The distribution of microbiome richness from individual samples across the two environments (sawfish; water). Richness is measured as the number of unique microbial species identified in the dataset. Circles represent samples sequenced using 2x150 bp inserts, while triangles represent 2x300 bp sequencing. The red dashed line corresponds to the mean richness across all samples (mean alpha diversity), and the blue dashed line corresponds to the gamma diversity, defined as the total number of unique species across each dataset. The size of the shape is representative of the library size.

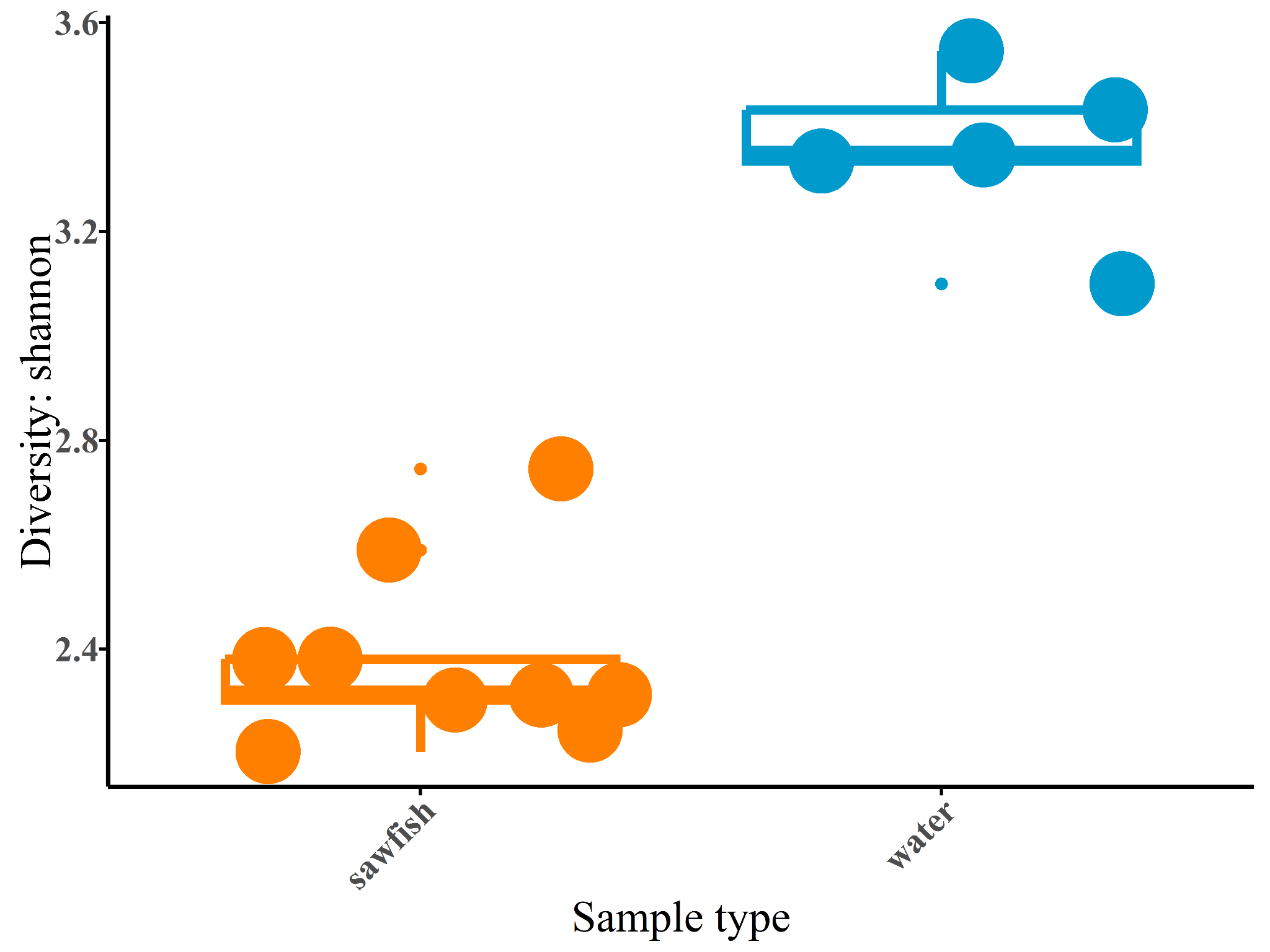

**SI Figure 4:** Boxplots showing family-level Shannon diversity of microbial communities associated with sawfish skin and the surrounding water column. Boxes represent the interquartile range with the median indicated by the central line; whiskers extend to 1.5 × the interquartile range, and points denote individual samples. Differences between host-associated and free-living communities highlight contrasts in diversity structure between the sawfish skin microbiome and the ambient water column.

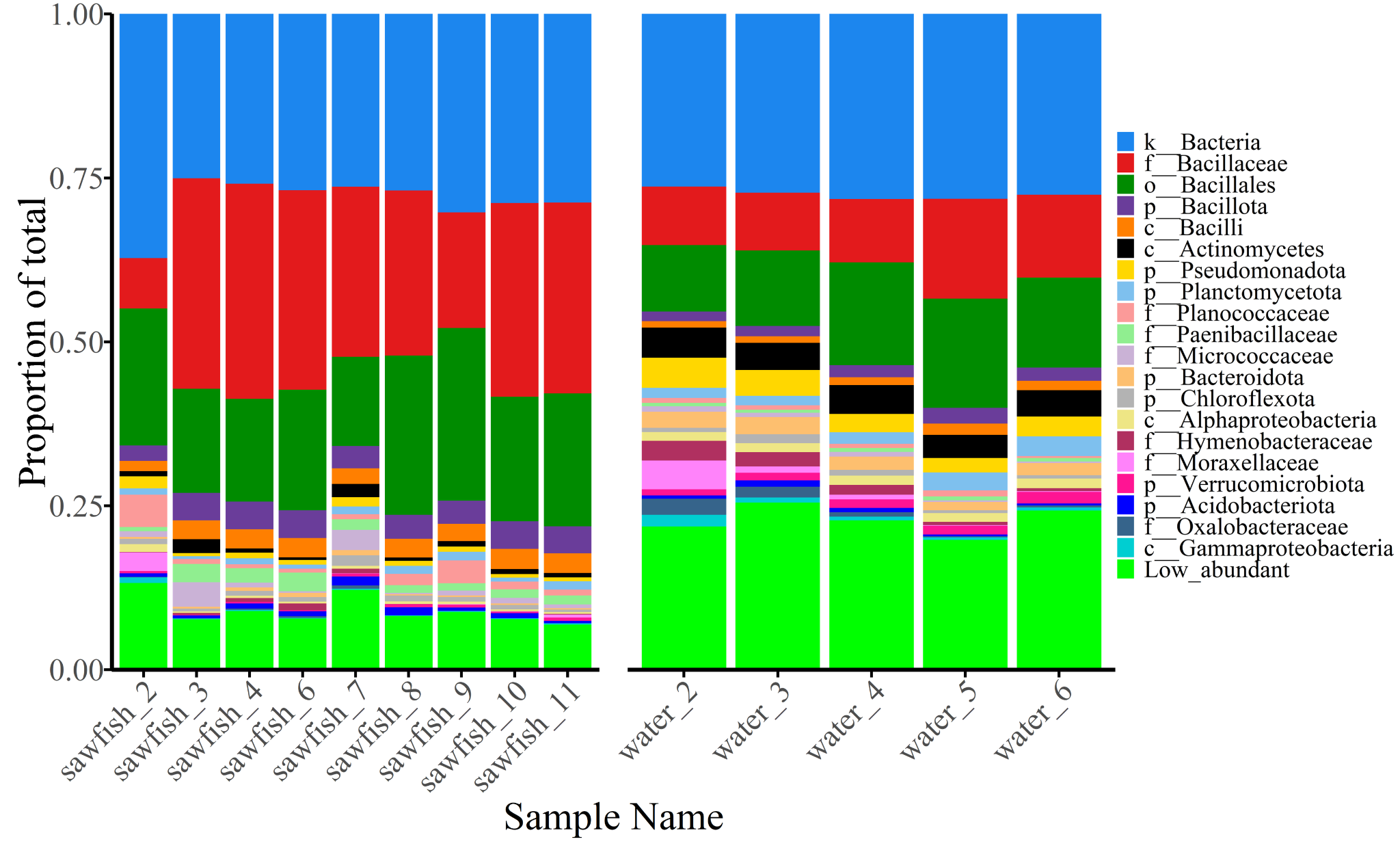

**SI Figure 5:** Microbial distribution of the top 20 microbial Families in sawfish and surrounding water column microbial communities at the taxonomic level of **f**amily. The x-axis are the samples from the sawfish and water. The y-axis represents the proportion of the total number of sequencing reads. The figure legend indicates the level which the microbial group is, for instance k__ indicates Kingdom, p__ Phylum, o__ Order, and f__ Family. Read sequences that were mapped equally across Families were assigned the common Order level, or Phylum. All other microbial Families are assigned to Low abundant.

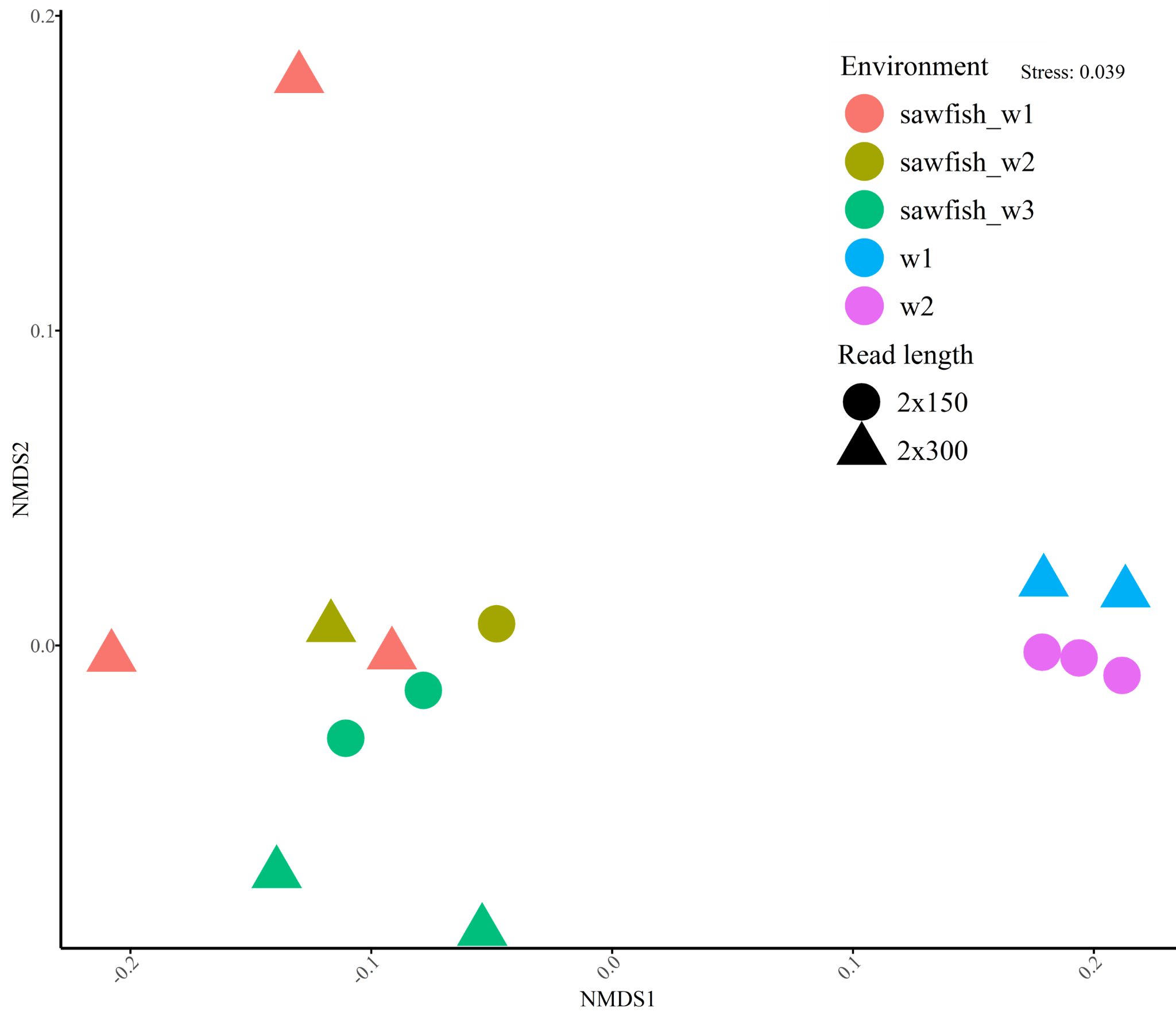

**SI Figure 6:** The microbial family level of the sawfish and water microbiomes represented by nMDS ordination plot. The distance between any two samples corresponds to the relative similarity in microbiome composition, measured by Bray-Curtis similarity from 4th-root transformed relative abundance data. Colour represents the day of sampling and the shape corresponds to the sequencing library size.

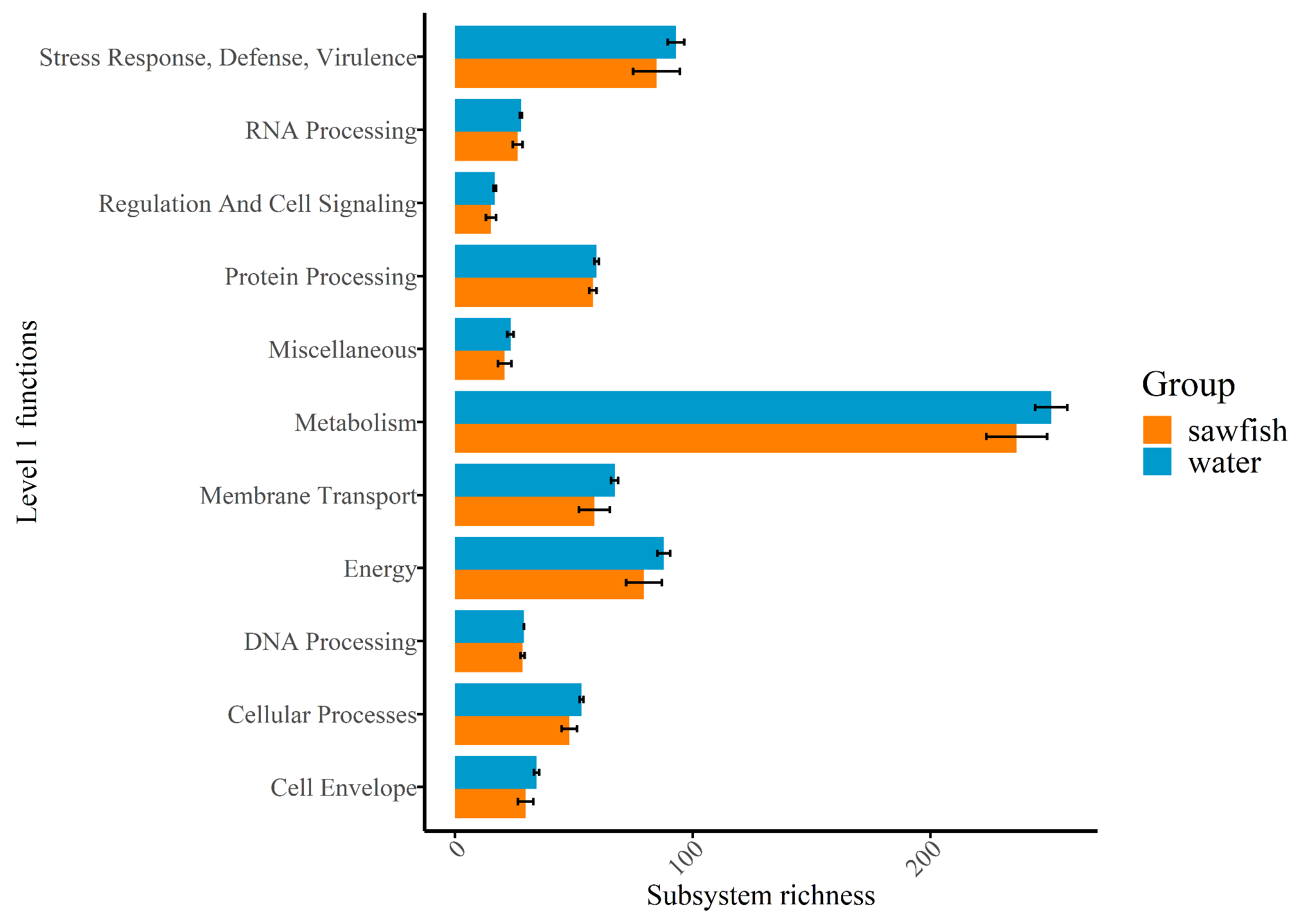

**SI Figure 7**: Richness of Subsystems across Level 1 SEED functional categories. Richness represents the number of Subsystems detected within each Level 1 category for sawfish and water column environments. Error bars indicate standard error. The x-axis shows the number of Subsystems, and the y-axis lists the Level 1 functional categories.

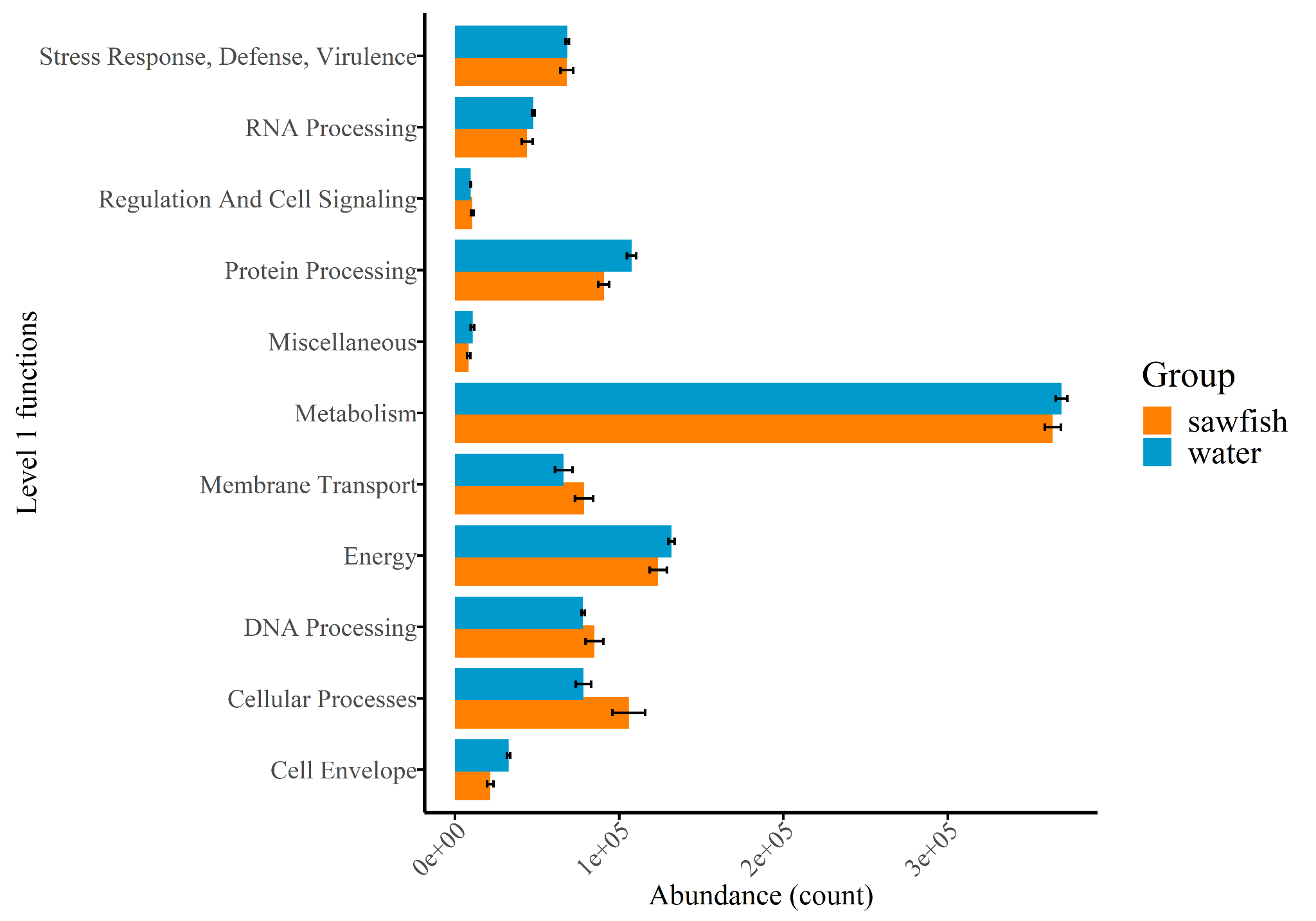

**SI Figure 8:** Abundance of Subsystems across Level 1 SEED functional categories. Abundance is represented as the number of sequencing reads assigned to each Subsystem within a Level 1 category for sawfish and water column environments. Error bars indicate standard error. The x-axis shows read count (abundance), and the y-axis lists the Level 1 functional categories.

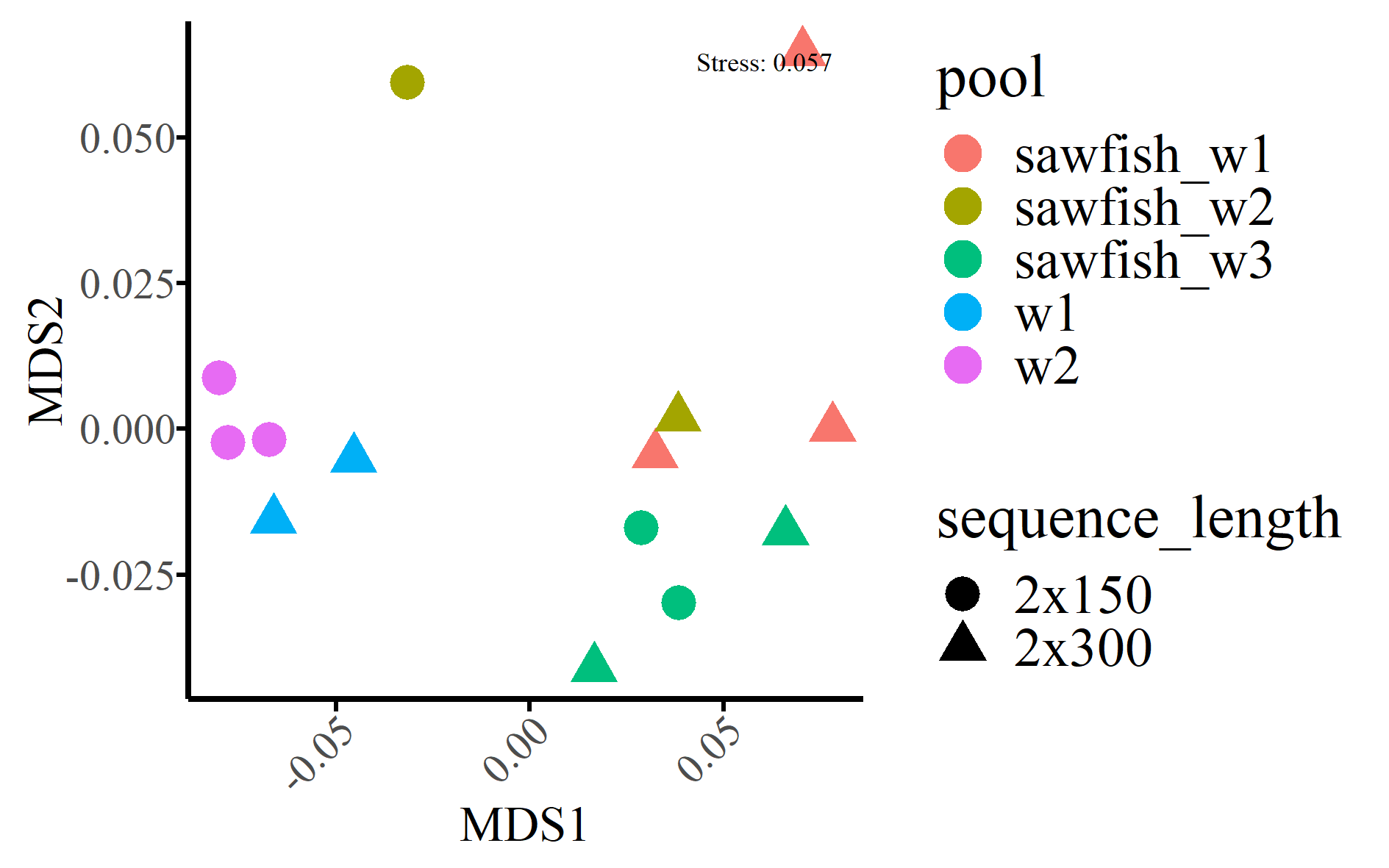

**SI Figure 9:** The gene functions of the sawfish and water microbiomes represented by nMDS ordination plot. The functions in sawfish and water microbiomes were distinctive, with high similarity among most replicate samples (samples from similar environments). Color represents the day and environment from which the sample was taken. Shape corresponds to the read length. The distance between any two samples corresponds to the relative similarity in microbiome composition, measured by Bray-Curtis similarity from 4th-root transformed relative abundance data.
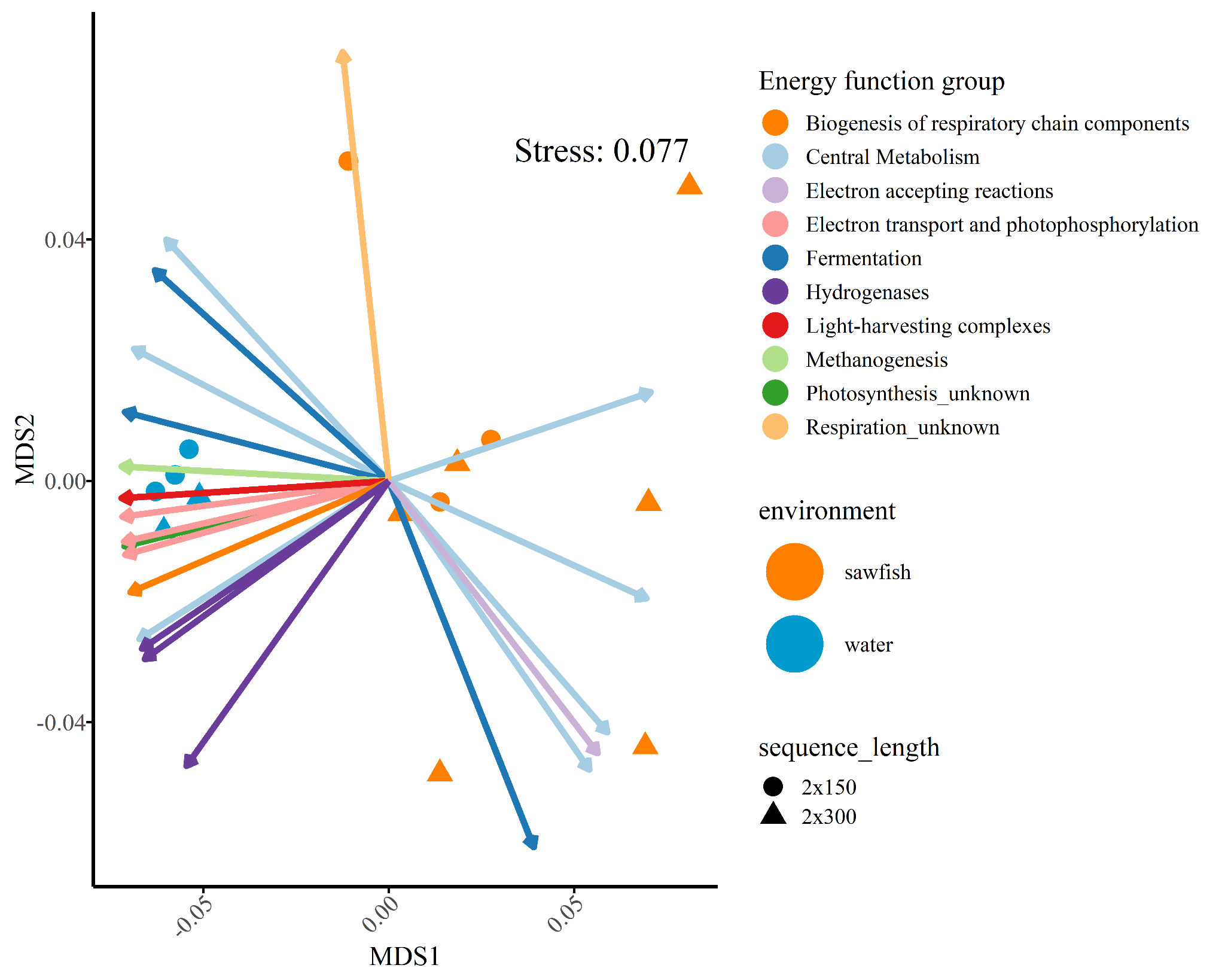

**SI Figure 10:** nMDS ordination of microbial energy subsystems from sawfish and water column environments. Circles and triangles represent metagenomic libraries of different sequence lengths, illustrating that environment is the primary driver of microbiome gene function patterns. Vectors indicate the direction of positive influence of each subsystem and are colored by their corresponding Level 3 category. Stress values indicate the goodness-of-fit of the nMDS ordination.

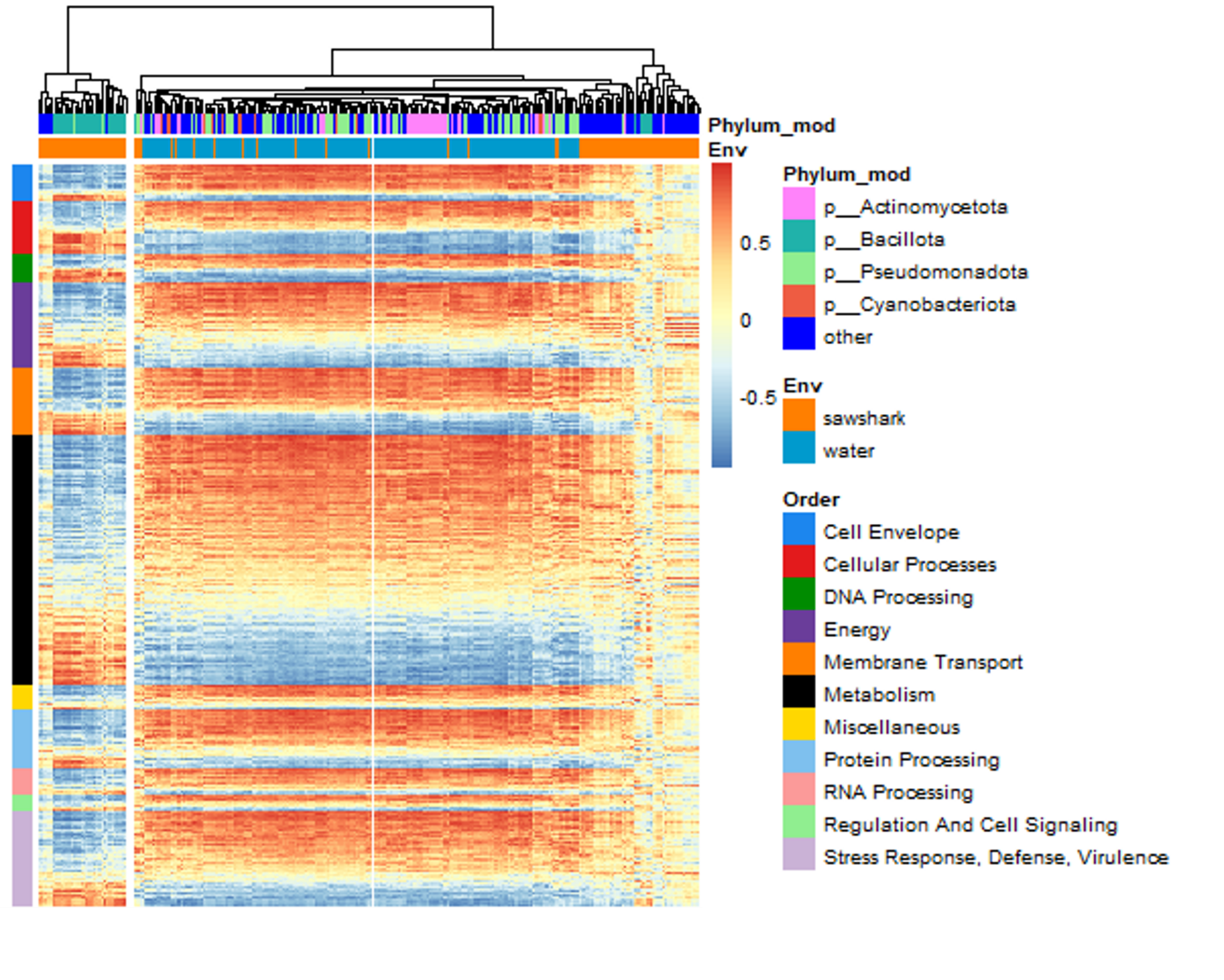

**SI Figure 11:** Heatmap showing the abundance relationship between all microbial taxa and gene functions from the sawfish and water column microbiomes. Red and blue correspond to positive and negative Spearman correlation, respectively. Lighter colors signify a weak relationship. On the x-axis, taxa are labelled by the phyla in which they are classified and the group from which they are significantly enriched. Gene functions are represented by rows and organized by their mean abundance across the dataset within the Level 1 SEED category (the broadest classification).
